# An arginine-rich nuclear localization signal (ArgiNLS) strategy for streamlined image segmentation of single-cells

**DOI:** 10.1101/2023.11.22.568319

**Authors:** Eric R. Szelenyi, Jovana S. Navarrete, Alexandria D. Murry, Yizhe Zhang, Kasey S. Girven, Lauren Kuo, Marcella M. Cline, Mollie X. Bernstein, Mariia Burdyniuk, Bryce Bowler, Nastacia L. Goodwin, Barbara Juarez, Larry S. Zweifel, Sam A. Golden

**Author notes:** Allen Institute for Cell Science, Seattle, WA, USA. University of Maryland School of Medicine, Department of Neurobiology, Baltimore, MD, USA. Co-corresponding authors: Eric R Szelenyi; Sam A. Golden, **Email:**.

## Abstract

High-throughput volumetric fluorescent microscopy pipelines can spatially integrate whole-brain structure and function at the foundational level of single-cells. However, conventional fluorescent protein (FP) modifications used to discriminate single-cells possess limited efficacy or are detrimental to cellular health. Here, we introduce a synthetic and non-deleterious nuclear localization signal (NLS) tag strategy, called ‘Arginine-rich NLS’ (ArgiNLS), that optimizes genetic labeling and downstream image segmentation of single-cells by restricting FP localization near-exclusively in the nucleus through a poly-arginine mechanism. A single N-terminal ArgiNLS tag provides modular nuclear restriction consistently across spectrally separate FP variants. ArgiNLS performance in vivo displays functional conservation across major cortical cell classes, and in response to both local and systemic brain wide AAV administration. Crucially, the high signal-to-noise ratio afforded by ArgiNLS enhances ML-automated segmentation of single-cells due to rapid classifier training and enrichment of labeled cell detection within 2D brain sections or 3D volumetric whole-brain image datasets, derived from both staining-amplified and native signal. This genetic strategy provides a simple and flexible basis for precise image segmentation of genetically labeled single-cells at scale and paired with behavioral procedures.

**Significance Statement:** Quantifying labeled cells in fluorescent microscopy is a fundamental aspect of modern biology. Critically, the use of short nuclear localization sequences (NLS) is a key genetic modification for discriminating single-cells labeled with fluorescent proteins (FPs). However, mainstay NLS approaches typically localize proteins to the nucleus with limited efficacy, while alternative non-NLS tag strategies can enhance efficacy at the cost of cellular health. Thus, quantitative cell counting using FP labels remains suboptimal or not compatible with health and behavior. Here, we present a novel genetic tagging strategy – named ArgiNLS – that flexibly and safely achieves FP nuclear restriction across the brain to facilitate machine learning-based segmentation of single-cells at scale, delivering a timely update to the behavioral neuroscientist’s toolkit.

## Introduction

The structure of the mammalian brain is complex and spans multiple spatial scales, requiring the development and application of spatially integrated methods to reveal key relationships with its function. High-throughput volumetric fluorescent microscopy pipelines, facilitated by light-sheet fluorescence microscopy (LSFM)^1^ and serial section block-face microscopy^2,3^, have accelerated these efforts. Notably, they have democratized experimental access to spatially integrated intact whole-brain anatomy, at the resolution of single-cells and within computationally feasible datasets that are compatible with brain atlas mapping. For example, in combination with genetic fluorescent labeling, these techniques have facilitated the mapping of cellular activity^4–16^, cell-type distributions^17–20^, and endogenous genetic^21^ and connectivity patterns^22^.

However, while significant hardware-based advancements have been made in microscopy systems^2,23,24^, and numerous software-based image analysis tools are readily available^5,13,14,25–29^, there has been a relative lack of genetic tool development that maximizes single-cell detection and quantification through the optimization of fluorescent signal. As opposed to hardware- and software-based approaches, genetic approaches may offer improved cellular detection without the prohibitively expensive acquisition of cutting-edge microscopy and computational equipment. Past genetic-based approaches have relied on fluorescent protein (FP) expression in its unmodified or modified forms^30^, but the efficacy of these prior efforts have not been systematically examined at the level of single-cell discrimination across image datasets of various sizes. As a result, the quantitative accuracy of single-cell brain mapping measurements remains ambiguous both within and between studies.

Typically, single-cell discrimination is improved by targeted genetic modifications to FPs by (*i*) N- or C-terminal tags of a minimal nuclear localization signal (NLS) sequence from a naturally occurring protein like the simian virus 40 large T antigen (SV40nls)^31–33^, or (*ii*) fusion with a full-length nuclear protein, like histone 2B^17,22,34^. Both approaches target FP localization to the cell nucleus to reduce extra-nuclear fluorescence. However, these approaches have limitations. Minimal NLS tag modifications only achieve partial nuclear localization, and therefore are sub-optimal for image segmentation. Conversely, full-length nuclear protein fusions guarantee nuclear restriction, but depend on overexpression of the fused nuclear gene, which can have significant deleterious effects on neuronal function and animal health^35^.

Two major bottlenecks have slowed the development of new ectopic NLSs - including those identified in silico^36^ or synthetic versions developed for genome-editing^37,38^ – for use as FP fusion tags. First, NLS performance when ectopically tagged as an FP fusion is unpredictable. This is because the nuclear import efficiency of naturally occurring NLSs are influenced by many factors, including intrinsic NLS sequence affinity to nuclear transport receptors KPNA and KPNB^39^, and their interplay with the amino acids surrounding the NLS in its native state^40,41^. Second, due to the barrier posed by challenging biological properties, there is not a high-throughput pipeline to systematically screen novel NLSs for whole brain mapping compatibility. Those properties include: (*i*) minimal extra-nuclear FP localization for high SNR, (*ii*) a simple and flexible sequence with minimal length and a single location of fused tag for unaltered FP function, (*iii*) modular compatibility with FP variants for spectral flexibility in imaging design, including multiplexing, (*iv*) equal functionality across cell-types for unbiased labeling of single-cells, and (*v*) non-deleterious effects of cellular and animal health. Thus, a genetic tag strategy that simultaneously synergizes and stereotypes FP-tagged NLS function while maintaining cellular health could satisfy these requirements.

Here, we present a new NLS genetic tag strategy named “Arginine-rich NLS” or “Argi-NLS”, that improves nuclear localization of genetically encoded FPs and enhances image segmentation of single-cells through a poly-arginine driven nucleolar enrichment mechanism. This method does not alter neural physiology or mouse behavior after standard AAV expression windows and is therefore compatible with AAV-paired behavioral procedures. It is functionally modular when using spectrally separate FPs and possesses equal efficacy when expressed within genetically distinct cortical cell-types. Moreover, it significantly improves two- and three-dimensional image segmentation that decreases computational expense associated with ML-based segmentation.

## Results

### Design of a nuclear-restricting Arginine-rich NLS (ArgiNLS)

Arginine (R) is the most hydrophilic, basic natural amino acid residue with a pKa over 12, and possession of a positive charge within all biological environments. Because of these properties, synthetic polyarginine (poly-R) tags (i.e., a peptide sequence consisting of multiple, R repeats) serve a variety of experimental functions, including their use as a protein purification tag^42–45^, cell-penetrating peptide signal, multi-DNA-binding domain linker, and nucleolar trafficking signal^42,46–48^. We hypothesized that a poly-R stretch coded in-between a NLS sequence and a downstream FP could act as an improved nuclear-localizing agent within neurons (**Fig. 1a**).

**Figure 1.**
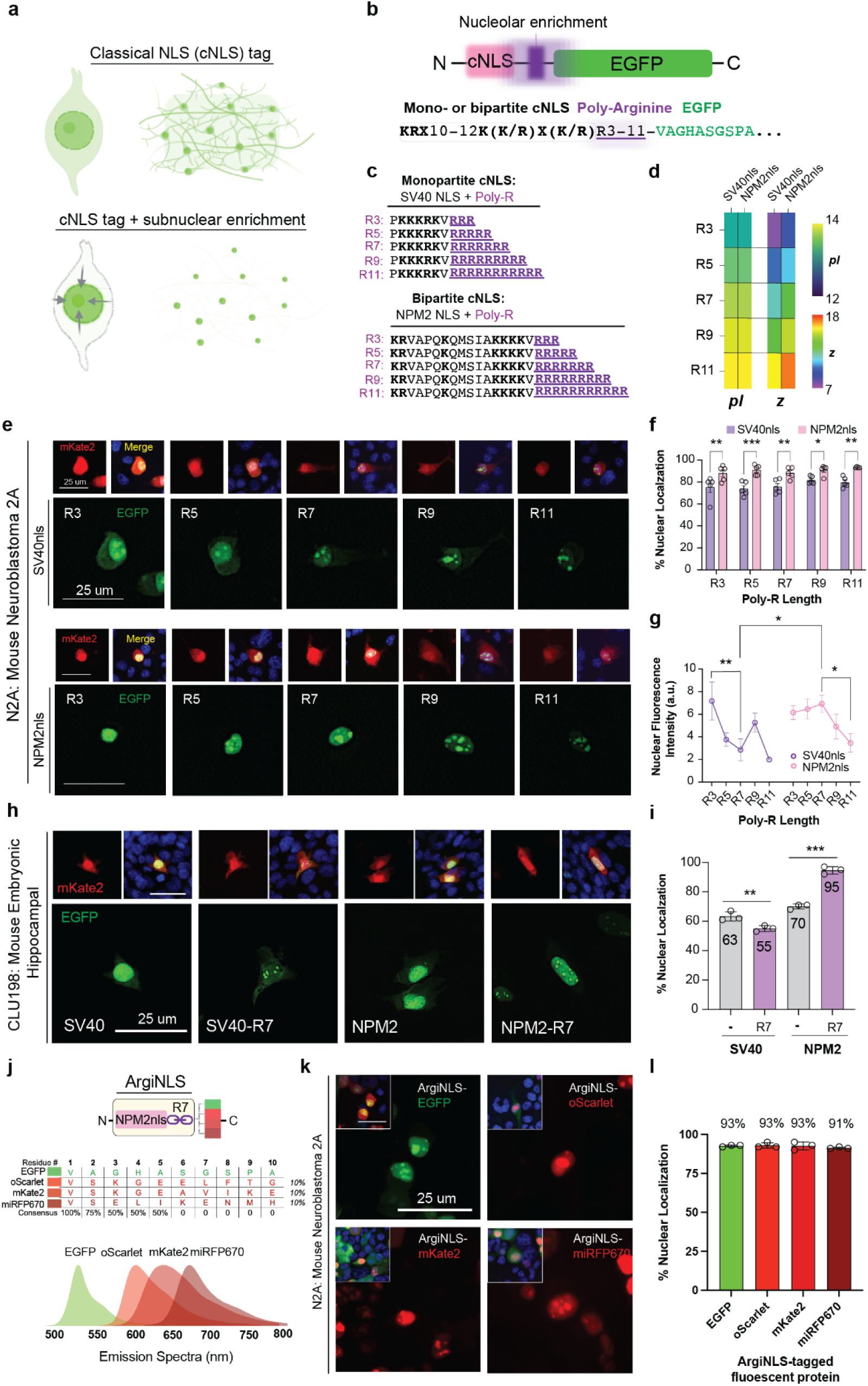
Strategy and *in vitro* characterization of an ariginine-rich NLS (ArgiNLS). **a**, Schematic depicting extracellular signal caused by classical NLS (cNLS) tag use, and depicting the theoretical effects of an optimized NLS tag that eliminates extra-nuclear signal artifact through a subnuclear enrichment strategy. **b**, Top: Configuration of cNLS, poly-arginine nucleolar enrichment tag, and EGFP in AAV overexpression vectors created for *in vitro* testing. Bottom: Corresponding amino acid composition of mono- or bipartite cNLS consensus sequences followed by poly-R stretch separating the coding sequence (CDS) of EGFP (first 10 amino acids shown). **c,** Amino acid sequences of SV40nls- and NPM2nls-poly-R tags. Basic amino acids are indicated in bold. **d**, Heat maps of isoelectric point (pI) and net positive charge (z) of each tag configuration. **e,** Representative images of transiently co-transfected single N2A cells displaying untagged mKate2 and SV40nls-poly-R (top) or NPM2nls-poly-R (bottom) tagged EGFP expression, spanning 3 to 11 R. **f,** Percent nuclear localization of each NLS across poly-R lengths (n=5/condition). **g**, Nuclear fluorescence intensity (a.u. = arbitrary units) across poly-R lengths for each NLS from same cells measured in **f**. **h**, Representative images of transient co-transfections with untagged mKate2 and each experimental EGFP-expressing construct within immortalized CLU198 embryonic hippocampal mouse cell line. **i,** Percent nuclear localization with/without R7 addition for each cNLS. **j**, Molecular schematic of ArgiNLS tag (NPM2nls-R7) and amino acid sequence comparison and consensus of the first 10 N-terminal residues to EGFP, oScarlet, mKate2, and miRFP670. **k**, Representative images of transient co-transfections with (from L to R and top to bottom) ArgiNLS-EGFP/mKate2, ArgiNLS-oScarlet/EGFP, ArgiNLS-mKate2/EGFP, and ArgiNLS-miRFP670/EGFP combinations within N2A mouse cell line. **l**, Percent nuclear localization of ArgiNLS-tagged FPs. **i** and **l** cell culture summary data is from 5 blinded and randomly selected cells quantified per transfection condition across 3 replicate cultures. The data in **f** was statistically analyzed using a two-way ANOVA followed by Holm-Sidak’s multiple comparison post hoc test. The data in **g** was statistically analyzed using a two-way ANOVA followed by Holm-Sidak’s multiple comparison post hoc tests for effects of poly-R length both within and across each cNLS. The data in **l** was statistically analyzed using a two-way (NLS (3) x +/- R7 (2)) ANOVA followed by Sidak post hoc test. *p<0.05; **p<0.01; ***p<0.005

To explore this idea in vitro, we first tested the effects of a range of poly-R lengths on the nuclear localization of two known classical NLS (cNLS) types: (*i*) monopartite (MP), composed of a single cluster of 4-8 basic amino acids and consensus recognition motif of K (K/R) X (K/R) (X is any amino acid), and (*ii*) bipartite (BP), composed of two clusters of 2-3 positively charged amino acids, separated by a 9-12 amino acid linker region, and with a consensus sequence R/K(X)10-12KRXX^39,49^ (**Fig. 1b; Table S1**). We selected cNLS sequences derived from the widely applied and characterized MP SV40^50,51^ or BP nucleoplasmin2 (NPM2)^52^ genes, whose NLS protein sequences differ in length and net positive charge but share the same isoelectric point (pI) across poly-R length additions (**Fig. 1c-d**). We performed co-transfections of AAV-(SV40nls or NPM2nls)-poly-R-EGFP overexpression constructs with untagged mKate2 in mouse neuroblastoma 2A (N2A) cells. Nuclear or cellular fluorescence localization and brightness (see **Methods** for a detailed description quantification approach) in individual EGFP+/mKate2+ cells were measured using DAPI staining as a nuclear marker and untagged mKate2 expression to label entire cells.

Co-transfected EGFP+/mKate+ cells revealed a poly-R length-dependent increase in nucleolar localization for both cNLSs. NPM2nls, and not SV40nls, exhibited decreased extra-nuclear fluorescence regardless of poly-R length (**Fig. 1e**). Measured composite values across poly-R lengths was 77% (± 1.48) for SV40nls and 91% (± 0.85) for NPM2nls (data not shown), where NPM2nls significantly obtained stable, higher nuclear localization across all poly-R lengths (**Fig. 1f**). Additionally, unlike the linear decrease observed with SV40nls, nuclear brightness with NPM2nls peaked with poly-R7 and decayed with longer lengths (**Fig. 1g**).

We validated the optimal NPM2nls-R7 configuration in a separate heterologous mouse brain cell-line by comparing untagged performance for each cNLS type (**Fig. 1h-i**). The control SV40nls tag exhibited 63% of nuclear EGFP fluorescence that was significantly decreased to 55% with the poly-R7 addition, despite a nucleolar-like localization (**Fig. 1h-i**). NPM2nls-R7 enhanced nuclear localization above untagged levels from 70 to 95% (**Fig. 1i**). We next probed the specificity of poly-R7 on BP cNLS activity using a BP version of SV40nls (biSV40nls, **Fig. S1**). We observed a significant enhancement of nuclear-localized EGFP fluorescence with biSV40nls-R7 (93% ± 1.5) versus untagged biSV40nls (70% ± 7.3) (**Fig. S1b-c**).

Prior work has shown that a synthetic poly-R8 tag facilitates DNA-binding protein function when inserted as a linker sequence^47^, suggesting that poly-R’s can be used as a linker in other molecular contexts. Next, we tested the ability of the NPM2nls-R7 genetic tag – now termed ArgiNLS – to provide modularity with other FPs beyond EGFP (**Fig. 1j-l**). We attached ArgiNLS to a range of spectrally separate FPs that have diverging N-terminal sequences located within the first 10 amino acids, which could influence fused NLS tag activity (**Fig. 1j**). Monomeric FPs included EGFP (Ex/Em: 488/507), oScarlet^53^ (Ex/Em: 569/594), mKate2^54^ (Ex/Em: 588/633), and miRFP670^55^ (Ex/Em: 647/670) (**Fig. 1j; bottom**). In N2A cells, we measured nuclear restriction of native fluorescence for EGFP (93% ± 0.3), oScarlet (93% ± 1.0), mKate2 (93% ± 1.5), and miRFP670 (91% ± 0.3) (**Fig. 1k-l**).

### Nuclear restriction of FP fluorescence *in vivo*

The *in vitro* results show enhanced ArgiNLS restriction of nuclear fluorescence. We next determined if the performance replicated *in vivo* using AAV overexpression in the mouse brain. Nuclear import of NLS-containing proteins – including SV40 and NPM2 - are controlled by nuclear transport receptors called importins (or karyopherins)^49^. Importins are dynamically expressed in multiple isoforms across cell-types and brain regions^56^, providing a potential cause of variable NLS activity of NLSs in a cell-type and brain region-dependent manner. We used a conditional AAV co-expression strategy to quantify nuclear-localized EGFP fluorescence in two genetically distinct cortical cell classes, GABAergic or glutamatergic neurons. We reasoned that each class comprises a divergent set of multiple neuronal cell sub-types that have functionally and genetically distinct characteristics^20,57^, and therefore are well suited for benchmarking the nuclear localizing efficacy of ArgiNLS- or conventional SV40nls-tagged EGFP.

We selected the barrel cortex – where GABAergic and glutamatergic cell-types are abundant^20,57^ – for AAV co-injections using SV40nls or ArgiNLS tagged EGFP (AAV-EF1a-SV40nls- *or* ArgiNLS-EGFP), combined with a conditional Cre-dependent virus to express somatic tdTomato (AAV-CAG-FLEX-tdTomato) in GABAergic or glutamatergic cells within *vGat-Cre* or *vGlut1-Cre* animals, respectively (**Fig. 2a**). Confocal images were acquired of the injection site, and nuclear (as defined by DAPI+ co-stain) and somatic (as defined by tdTomato+ somatic fluorescence) EGFP fluorescence was quantified at the level of single-cells across NLS tags and cell-types at both high (**Fig. 2b-c**) and low (**Fig. S2**) magnification.

**Figure 2.**
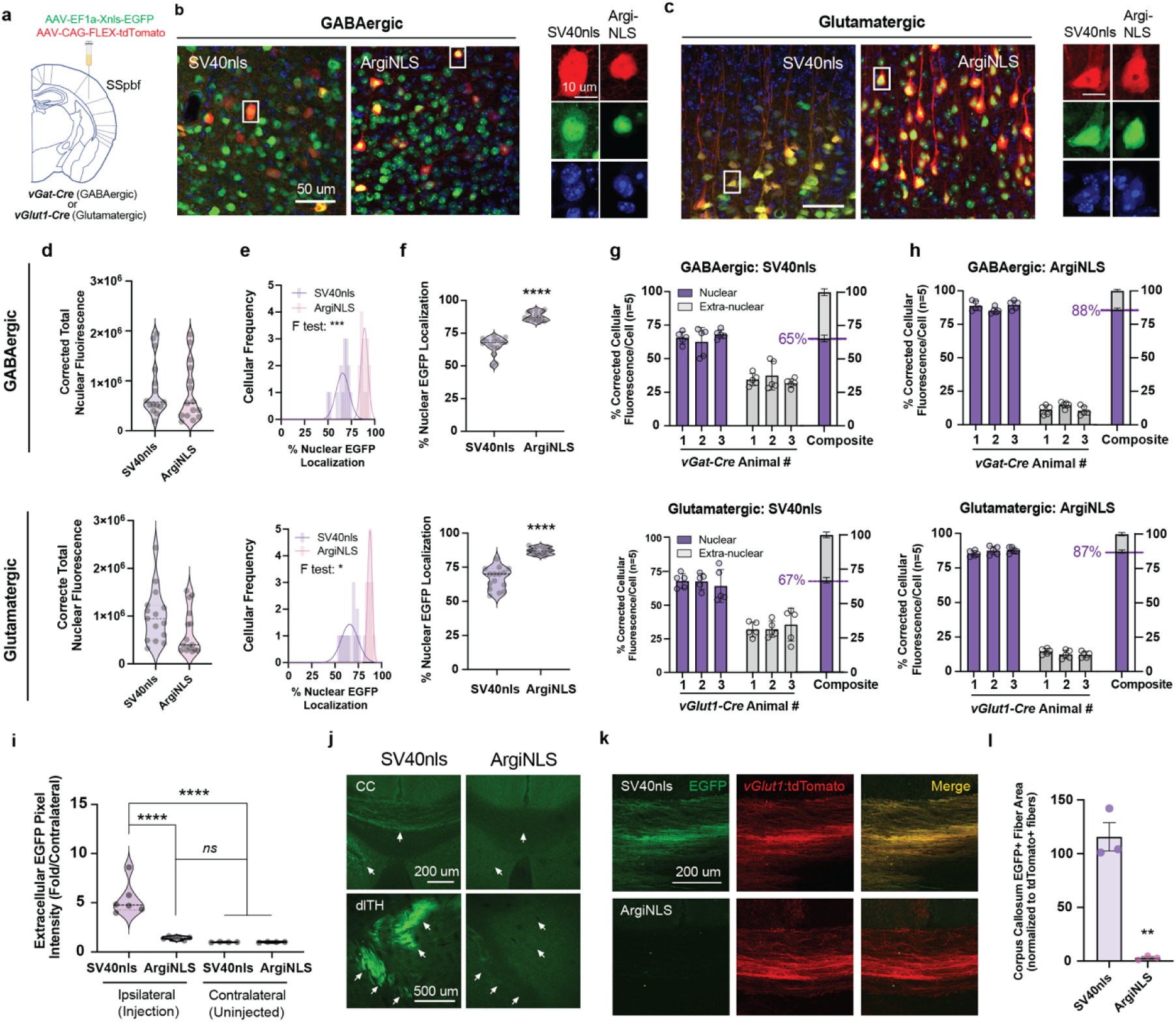
Optimized nuclear localization of fluorescence across major cortical cell-type classes in vivo. **a,** Schematic of stereotaxic AAV co-injection in the barrel field of primary somatosensory cortex (SSpBF) of *vGat-Cre* or *vGlut1-Cre* transgenic mice. **b**-**c**, Representative 40x confocal images of SV40nls-EGFP/FLEX-tdTomato and ArgiNLS-EGFP/ FLEX-tdTomato viral expression in (**b**) *vGat-Cre* or (**c**) *vGlut1-Cre* mice. Right: Cropped representative image of an example EGFP+/tdTomato+ cell. **d-h,** Quantification of NLS tag performance in GABAergic (top) or glutamatergic (bottom) EGFP+/tdTomato+ cells (n=random 5 cells/animal; n=3 animals per Cre-driver/tag). **d**, EGFP corrected total nuclear fluorescence (CTNF). **e**, Cellular frequency histogram of percent nuclear EGFP localization. **f**, Percent nuclear EGFP fluorescence comparison across tags. **g-h**, Nuclear and extra-nuclear EGFP fluorescence for all quantified cells of each animal across (**g**) SV40nls- or (**h**) ArgiNLS-labeled cells. The composite percent nuclear EGFP across all animals is indicated to the right. **i**, Total pixel intensity of extracellular (EC) EGFP signal ipsilateral to injection site, expressed as a fold change from non-injected contralateral side (n=3 EC quantifications/3 *vGlut1-Cre* mice**). j**, Representative images of extranuclear EGFP signal in axon bundles of passage in the corpus callosum (CC; top) or terminations in the dorsal thalamus (dTH; bottom). Arrows indicate location of labeling observed with SV40nls and are superimposed on ArgiNLS-EGFP. **k**, Representative 40x confocal images of SV40nls- or ArgiNLS-EGFP with *vGlut1:*tdTomato viral expression in fibers of the CC (n=3 EC quantifications/3 *vGlut1-Cre* mice). **l**, Quantification of EGFP+ fiber area in CC normalized to *vGlut1*:tdTomato+ fiber area. All bar graphs represent mean ± SEM. The data in **d, f,** and **l** was statistically analyzed using an unpaired t-test. Group differences in population variance of data in **e** were statistically compared using an F-test. The data in **i** was statistically analyzed using a 1-way ANOVA followed by Tukey’s multiple comparison post hoc test. Violin plots display median and interquartile range as dashed lines. *p<0.01; **p<0.05; **p<0.01; ****p<0.001

Barrel cortex neurons infected with SV40nls-EGFP exhibited somatic EGFP concomitant with high extracellular background, putatively from SV40nls-EGFP localized in cellular processes (**Fig. 2b-c; left**). Conversely, ArgiNLS-EGFP did not exhibit extra-cellular EGFP background, which corresponded with nuclear-restricted EGFP and localization in presumptive nucleolar puncta (**Fig. 2b-c; right**). Sampled EGFP+/tdTomato+ single-cells used for nuclear/somatic EGFP fluorescence quantifications contained equal distributions of nuclear localization (corrected total nuclear fluorescence, CTNF; see **Methods**) levels across tags and cell-type (**Fig. 2d**). However, when CTNF was expressed as a percent cellular fluorescence (corrected total cellular fluorescence, CTCF; see **Methods**), nuclear localization variance was significantly different across tags; SV40nls showed high variance while ArgiNLS showed low variance (**Fig. 2e**). Despite differences in variance, tag type had a significant effect on nuclear localization, where ArgiNLS-EGFP contained significantly higher nuclear localization (GABAergic cells: 88% ± 0.9; glutamatergic cells: 87% ± 0.7) than SV40nls-EGFP (GABAergic cells: 65% ± 1.8; glutamatergic cells: 67% ± 2.0) (**Fig. 2f-h**).

We next determined how ArgiNLS impacts extracellular (EC) background fluorescence at the site of injection and within anatomically distal areas (**Fig. 2i-l**). First, we compared EC pixel intensity measurements taken from field of views (FOVs) positioned in-between EGFP-labeled cells of the injected barrel cortex (i.e., ipsilateral), and in-between DAPI+ cells of the uninjected (i.e., contralateral) barrel cortex, as a control measurement, from the same mice (**Fig. 2i**). Ipsilateral SV40nls FOVs contained a significant 4-fold increase in EC fluorescence compared to uninjected contralateral FOVs, that was also significantly higher than between-subject ipsilateral and contralateral ArgiNLS FOVs. EC fluorescence within ipsilateral ArgiNLS FOVs was unchanged from contralateral FOV levels from both SV40nls and ArgiNLS injected samples.

Neurons within the barrel cortex send diverse axonal projections throughout the brain^58^. ArgiNLS consistently eliminated non-nuclear labeling of axon fiber of passage in the corpus callosum (CC), as well as axonal terminations in the dorsolateral thalamus (dlTH), that was observed with SV40nls-EGFP injections (**Fig. 2j**). We then confirmed decreased CC axon labeling by ArgiNLS in separate quantifications made of EGFP sampled in high-magnification FOVs. To control for error in sample-to-sample variability of viral delivery, we restricted our analysis within the *vGlut1-Cre* injected sections and normalized EGFP+ fiber area to *vGlut1*:tdTomato+ fiber area (**Fig. 2k-l**). Accordingly, SV40nls-EGFP labeled 115% (± 13) of *vGlut1*:tdTomato+ fibers, indicating that extra-nuclear EGFP localization produced from constitutively expressed SV40nls-EGFP labels CC fibers within both *vGlut1(+*) and *vGlut1(-)* neurons of the barrel cortex (**Fig. 2l**). Conversely, ArgiNLS-EGFP labeled 3% (± 1.2) of *vGlut1*:tdTomato+ fibers.

### Non-deleterious to neuronal and behavioral health

Full-length nuclear protein-FP fusions, such as H2B-EGFP^59^, demonstrate high levels of nuclear-restricted fluorescence, but cause detrimental effects to cellular and animal health when overexpressed in the mouse brain^35^. Since ArgiNLS tagged FPs display similar nuclear-restricted fluorescence, we next determined if ArgiNLS-EGFP viral overexpression in the brain causes detrimental effects to baseline neuronal physiology or animal behavior (despite it lacking the fusion of an endogenous nuclear protein) (**Fig. 3**). We first examined baseline physiological properties of glutamatergic neurons within the barrel cortex of *vGlut1*-*Cre* animals. We used a local unilateral injection of Cre-dependent tdTomato virus to create control *vGlut1-Cre*:tdTomato(+) labeled neurons, and a contralateral co-injection of Cre-dependent tdTomato with ArgiNLS-EGFP viruses to create within-sample, experimental ArgiNLS-EGFP(+)/*vGlut1-Cre*:tdTomato(+) labeled neurons (**Fig. 3a**). Analysis of baseline physiological parameters indicated no effect of ArgiNLS-EGFP overexpression on resting membrane potential (**Fig. 3b**) and action potential generation across a range of injected currents (**Fig. 3c-d**).

**Figure 3.**
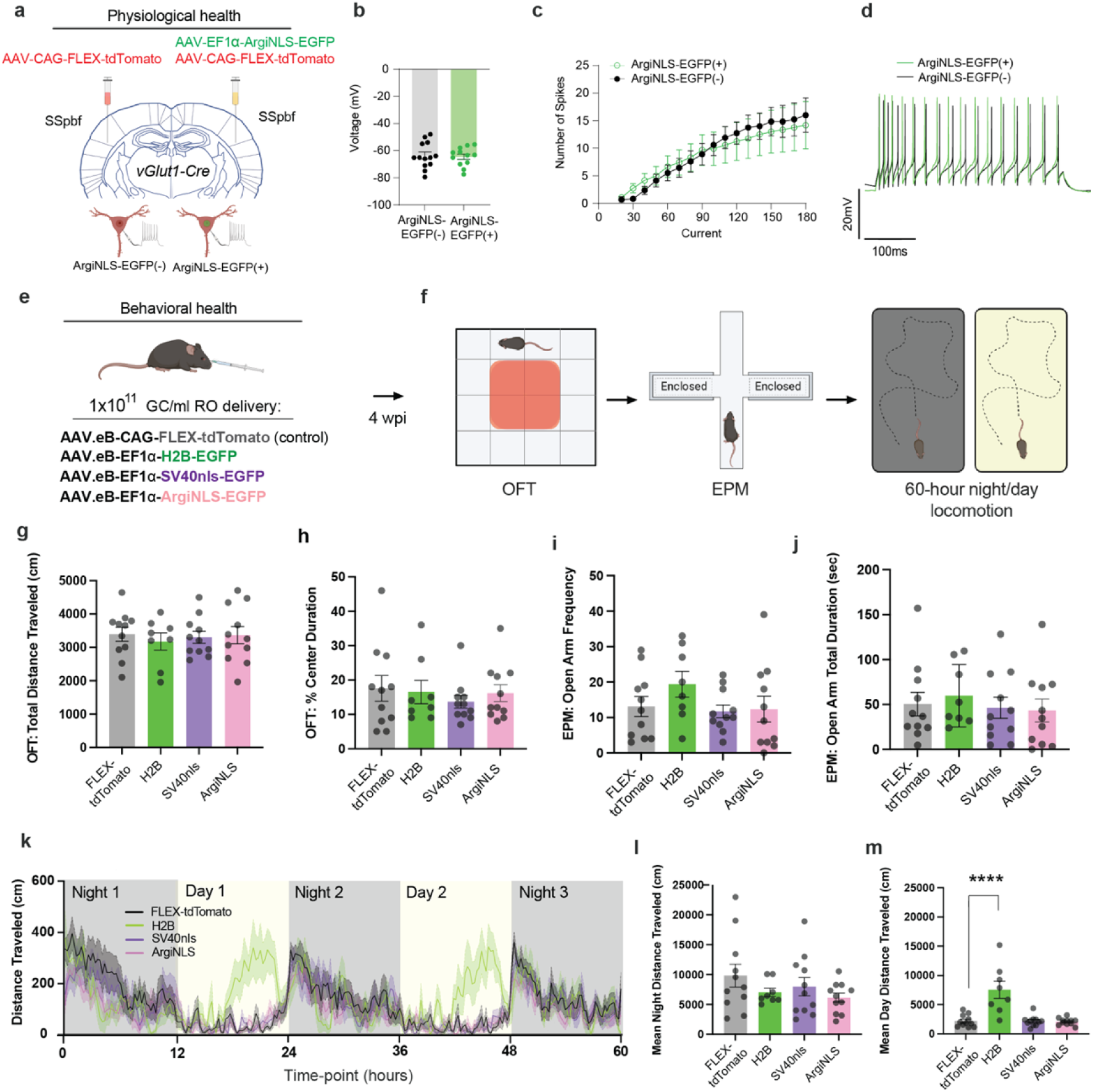
ArgiNLS-EGFP viral expression is not associated with physiological or behavioral deficits. **a,** Schematic illustration of bilateral stereotaxic AAV injection/co-injection in barrel cortices (SSpBF) of *vGlut1-Cre* transgenic mice, and electrophysiological recordings conducted on resulting *vGlut1-Cre*:tdTomato(+) neurons that are either ArgiNLS-EGFP(-) or ArgiNLS-EGFP(+). **b**, Resting membrane potential of ArgiNLS-EGFP(-) (n=13) or ArgiNLS-EGFP(+) (n=12) neurons. **c**, Current injection curve from ArgiNLS-EGFP(-) or ArgiNLS-EGFP(+) neurons. **d**, Example trace current injection at 100 pA. **e,** AAV.eB delivery and vector information schematic. Groups of mice were injected with 1 x 10^11^ GC of a negative control AAV.eB-CAG-FLEX-tdTomato (n=11), AAV.eB-EF1a-H2B-EGFP (n=8), AAV.eB-EF1a-SV40nls-EGFP (n=11), or AAV.eB-EF1a-ArgiNLS-EGFP (n=11). **f,** Baseline behavioral battery 4 weeks post viral injection (wpi). Mice were sequentially tested in the open field test (OFT), elevated plus maze (EPM), and 60-hour night/dark locomotion recording. **g**, Total distance traveled in the OFT. **h**, Percent time duration spent in the center of the OFT arena. **i**, Frequency of entries in the open arm of the EPM. **j**, Total duration of time spent in the open arm of the EPM. **k**, Locomotion XY plot showing total distance traveled across time of the 60-h recording. Day (yellow)/night (gray) phase of light cycle is underlaid the plot according to time of 60-h recording. **l**, Mean night distance traveled across the 60-h recording period. **m**, Mean day distance traveled across the 60-h recording period. All bar graphs represent mean ± SEM. The data for each panel from **g-j** and **l-m** were statistically analyzed using an one-way ANOVA with Dunnet’s multiple comparison posthoc test of each experimental group versus control FLEX-tdTomato group. ****p<0.0001

Next, we used the PHP.eB viral serotype to examine the effect of systemic whole-brain ArgiNLS-EGFP or control virus expression – including a non-expressing negative Cre-dependent tdTomato control virus, and positive SV40nls-EGFP and H2B-EGFP expressing controls - on behaviors including the open field task (OFT) and elevated plus maze (EPM) to study effects on acute exploratory drive, and a 60-hour locomotion monitoring assay to study chronic locomotion and sensorimotor performance across dark/light cycles (**Fig. 3g-m**). We found no differences in acute OFT performance across viral groups as measured by mean total distance (cm) traveled (**Fig. 3g**), or percent center time duration (**Fig. 3h**). Additionally, acute EPM performance as measured by open arm entry frequency (**Fig. 3i**), or open arm total duration (sec) (**Fig. 3j**) was the same across viral groups. Consecutive monitoring of homecage distance traveled (cm) across 3 nights and 2 days indicated a lack of effect from ArgiNLS-EGFP or SV40nls-EGFP systemic overexpression (**Fig. 3k-m**). Conversely, H2B-EGFP-injected animals displayed dysregulated locomotor activity during the light phase. We confirmed these behavioral data using 2D histology and imaging of native fluorescence on a subset of animals (**Fig. S3**).

### Enhanced compatibility with machine learning classification

Image classification of objects can be reliably automated and scaled through the creation of machine learning (ML) algorithms or “classifiers” that rapidly extract features based on pixel content to create image segmentations for quantification^60,61^. For fluorescently labeled images, the manual effort taken to compile training datasets is directly tied to the fluorescent signal’s fidelity to label a target class. Thus, poorly defined labeling may require extensive training and user annotation. Since ArgiNLS enables consistent FP restriction in the nucleus of neurons, we next asked if this influences the creation – or performance - of a single-cell ML classifier, as compared to a classifier trained with cSV40nls tag labeling (**Fig. 4**).

**Figure 4.**
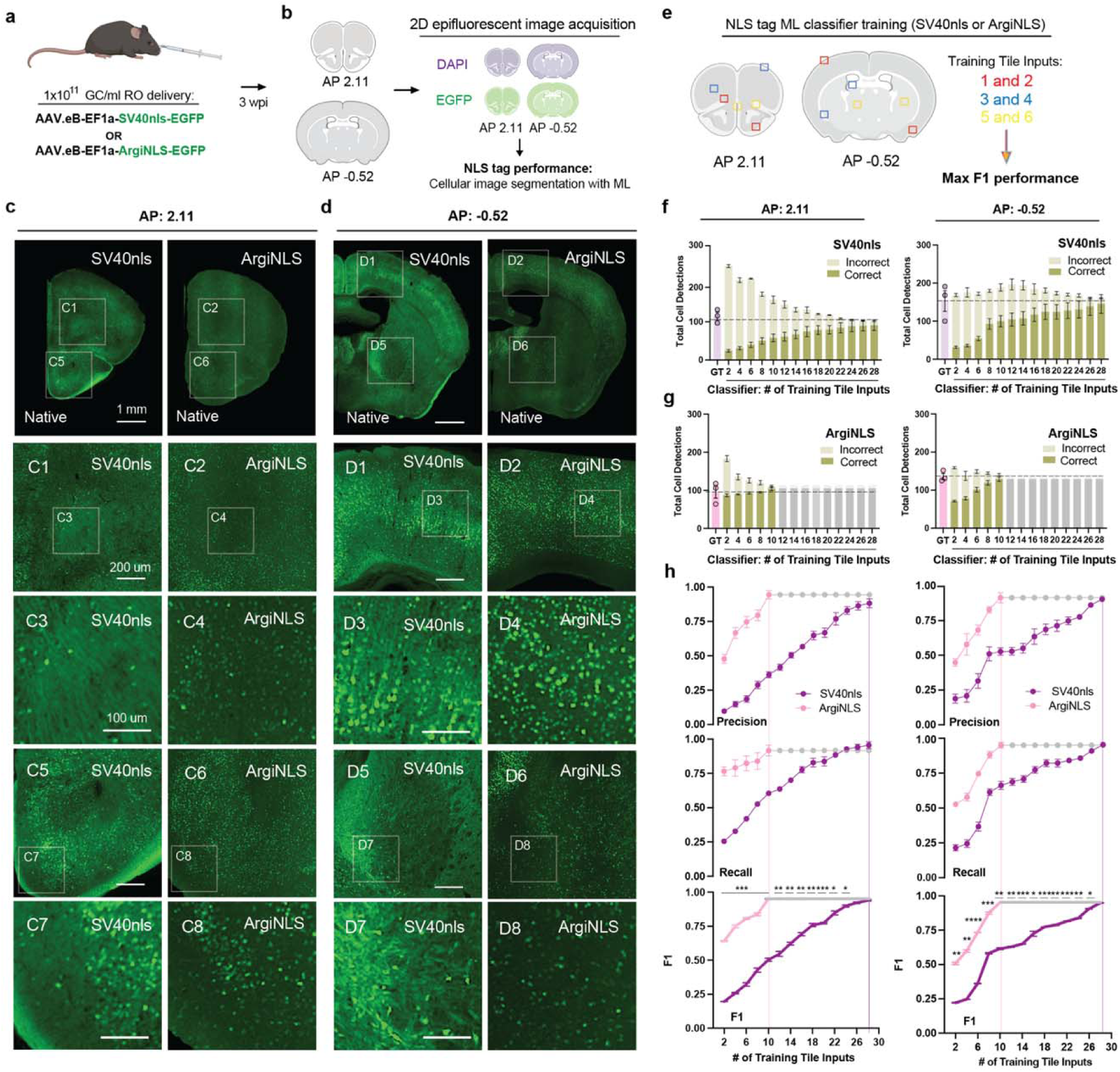
Enhanced performance of single-cell ML classification. **a**, Experimental schematic of systemic AAV.eB delivery of NLS-tagged EGFP. **b**, Microscopy acquisition schematic. **c-d**, Representative images of EGFP viral expression at matching (**c**) AP 2.11 or (**d**) −0.52 hemispheric coronal planes, across 2 FOVs at 3 magnification levels. **e**, Schematic of SV40nls and ArgiNLS classifier training workflow for single-cell segmentation of 2D images. **f**-**h**, Cell segmentation classifier benchmarking and performance metrics for (left) AP: 2.11 and (right) AP: −0.52 brain sections. Total cell detections for (**f**) SV40nls-EGFP or (**g**) ArgiNLS-EGFP for expert rater ground truth (GT; left bar) (n=3) and ML classifier across iterations of 2-increment additive training input rounds. Gray bars represent iterations after peak performance criteria was met. Dashed line indicates mean GT cell detections. **h**, Precision, recall, and F1 harmonic scoring of single-cell detection for SV40nls-EGFP and ArgiNLS-EGFP. Gray data points represent iterations after max F1 performance was met. All bar graphs represent mean ± SEM. The F1 data in **h** was statistically analyzed using a two-way repeated measures ANOVA followed by Holm-Sidak’s post hoc test. *p<0.05; **p<0.01; **p<0.01; ****p<0.001

We systemically labeled single-cells with SV40nls-EGFP or ArgiNLS-EGFP-expressing AAV.eB viruses through RO injections (**Fig. 4a**) and used DAPI counterstained images of coronal sections (2.11 and −0.52 from bregma) to benchmark single-cell ML classifier training and testing at varying anatomical locations throughout the brain (**Fig. 4b**). We observed that SV40nls displayed variable extra-nuclear fluorescence and background, and ArgiNLS conversely showed nuclear-restriction and an overall higher density of labeled single-cells (**Fig. 4c-d**).

We next created single-cell classifiers for SV40nls and ArgiNLS labels and benchmarked their performance using F1 scores: a machine learning metric that determines model accuracy based on a combination of precision (i.e., positive predictions) and recall (i.e., correct identification of positive class) measurements. We performed multiple training iterations consisting of two sample FOVs per session until maximum classifier performance, as measured by F1 scoring in FOVs with similar ground truth (GT) cell detections, was reached (**Fig. 4e**; **Fig. S4**). ArgiNLS (5 rounds, 10 input FOV) resulted in a substantial reduction in the total number of training iterations required to reach max classifier performance at both anatomical planes compared to SV40nls (14 rounds, 28 input FOV) (**Fig. 4f-h**). Specifically, for AP 2.11, SV40nls classification started with a 93% greater total amount of cell detections than GT, that was composed of 11% correct (24 ± 3.1) and 89% (222 ± 3.8) incorrect detections. Both correct and incorrect detections displayed an exponential-like scaling towards GT levels as function of training round number (**Fig. 4f; left**). Initial ArgiNLS classification consisted of 63% more total cell detections (184 ± 7.5; Correct: 87 ± 7.0; Incorrect: 96 ± 13.9) than GT, with correct cells matching 90% of GT (**Fig. 4g; left**), and incorrect cells progressively reduced over each training session.

Total cell detections for both classifiers at AP −0.52 was stable across all training iterations and corresponded closely to GT detection levels with training (**Fig. 4f-g; right**). Like the raw data, ArgiNLS classification outperformed SV40nls classification in both precision (**Fig. 4h, top**) and recall (**Fig. 4h, middle**). ArgiNLS significantly outperformed SV40nls in F-sore across nearly all training tile input FOV iterations (**Fig. 4h; bottom**).

### Enriched brain-wide segmentation of single-cells

We next sought to determine ArgiNLS tag’s impact on single-cell segmentation within a whole-brain mapping dataset using large-scale ML classification. We systemically delivered PHP.eB SV40nls-EGFP and ArgiNLS-EGFP expressing viruses and performed iDISCO+ whole-mount clearing and EGFP staining to fluorescently label all the infected cells throughout the brain (**Fig. 5a, top; 5b**). Whole-brain samples were imaged with light sheet fluorescent microscopy (LSFM), and image datasets were run through a pipeline consisting of reference atlas registration, ML cellular segmentation, atlas segmentation of brain regions, and voxelized cell density analyses (**Fig. 5a, bottom**). Like our observations in 2D histology, SV40nls-EGFP staining produced higher image artifact and less cells counts than ArgiNLS-EGFP (**Fig. 5b, left**). We therefore trained separate ML classifiers using ilastik software to achieve optimal single-cell classification performance, separately for each tag (**Fig. S5a**). ArgiNLS reduced the model training requirements in 3D, where SV40nls classifier training required a total dataset of 44 training images and ArgiNLS required 20 training images (**Fig. S5b**).

**Figure 5.**
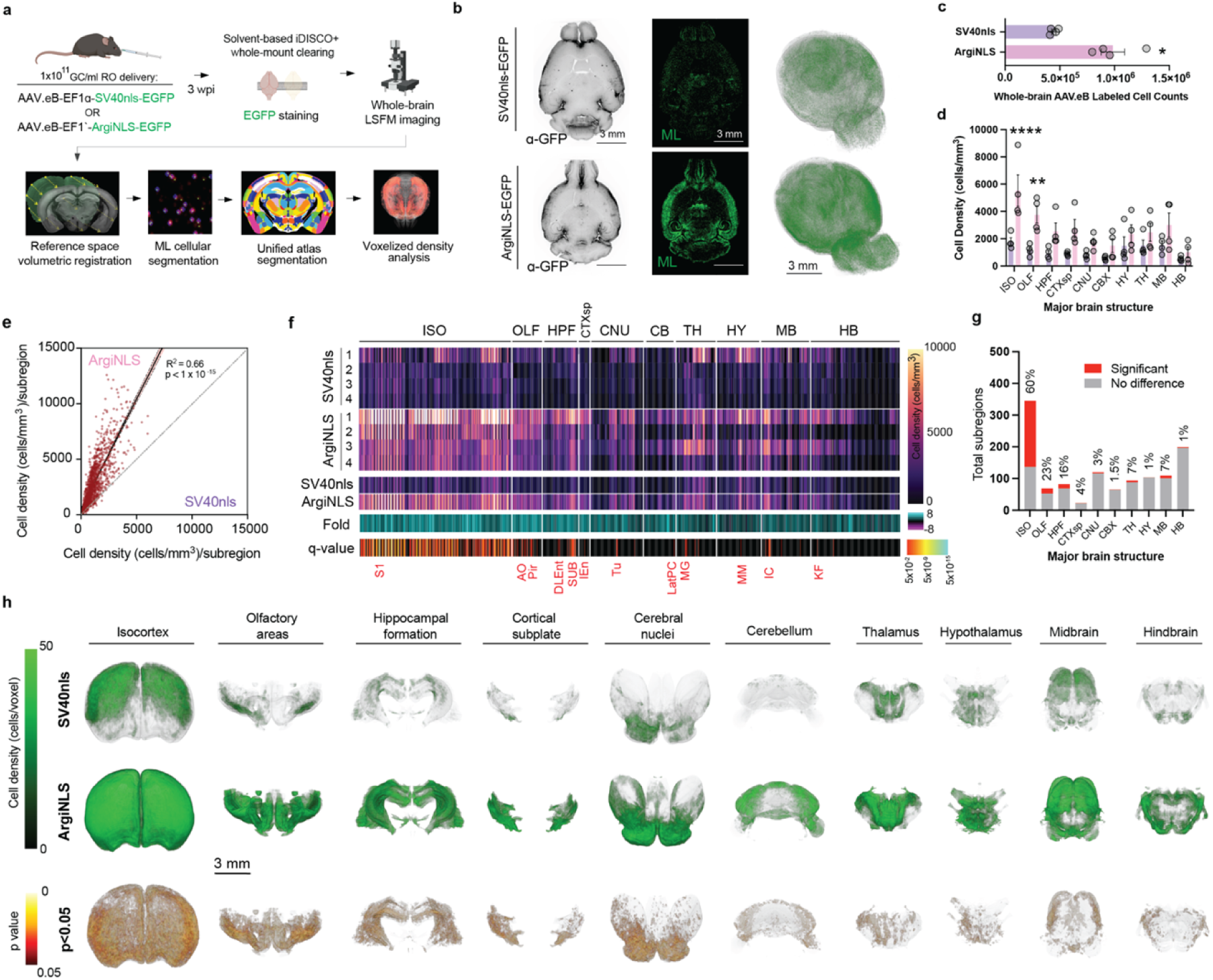
Enriched brain-wide segmentation of single-cells. **a**, Schematic of experimental and image processing workflow used to quantify systemic SV40nls- or ArgiNLS-EGFP-labeled cells across the whole mouse brain. **b,** Left: Raw LSFM image in horizontal orientation from a representative SV40nls-EGFP (top) or ArgiNLS-EGFP (bottom) expressing brain sample. Middle: corresponding ML-segmented cells within the same optical section. Right: Whole-brain 3D-renderings of voxelized cell density for each sample. **c,** Comparison of whole-brain AAV.eB-labeled cell counts. **d,** Comparison of cell density across major ontological brain structures. **e**, Scatterplot and linear regression of ArgiNLS (Y-axis) versus SV40nls (X-axis)-labeled mean cell density across all subdivisions of the brain. **f**, 2D heatmap plot of (from top to bottom): cell density across each individual sample, mean cell density, fold change over ArgiNLS, and statistical significance of mean cell density differences across all anatomical subdivisions by structure order. Major ontological brain structure positioning is indicated with white lines and written above. Example statistically significant subdivisions are listed in red. **g,** Bar graph of total statistically significant subdivisions versus no difference plotted for each major brain structure. Percent significant subregions is listed above each bar. **h**, Volumetric renderings of mean SV40nls-EGFP or ArgiNLS-EGFP voxelized cell densities (top) and the statistically significant voxels of the mean differences (bottom) across all major brain structures. All bar graphs represent mean ± SEM. The data for panel **c** was statistically analyzed using an unpaired Student’s t-test. The data for panel **d** was statistically analyzed using a two-way ANOVA with Bonferroni post-hoc test for mean significant differences at each major region. The data for panel **e** was analyzed with linear regression. The data for panel **f** was analyzed with FDR-corrected multiple t-tests using the two-stage step-up method of Benjamini, Krieger, and Yekutieli. *p<0.05; **p<0.01. Acronyms: ISO = isocortex; OLF = olfactory areas; HPF = hippocampal formation; CTXsp = cortical subplate; CNU = cerebral nuclei; CBX = cerebellum; HY = hypothalamus; TH = thalamus; MB = midbrain; HB = hindbrain; AO = accessory olfactory area; Pir: piriform cortex; DLEnt: dorsolateral entorhinal cortex; SUB: subiculum; Tu = olfactory tubercle; MG = medial geniculate; MM = medial mammillary area; IC = inferior colliculus; KF = Koelliker-Fuse subnucleus

Following extensive classifier validation (**SI Text**), we applied our whole-brain image processing pipeline to create maps of systemic AAV.eB infection. We found that ArgiNLS-EGFP significantly increases AAV.eB labeled whole-brain cell counts by 2.6-fold (SV40nls: 514,422 ± 90,388; ArgiNLS: 1,362,466 ± 268,889) (**Fig. 5c**). Additionally, we asked if the whole-brain count difference was anatomically distributed preferentially or evenly across gross areas of the brain. Statistically significant increases of ArgiNLS-labeled cell densities were found specifically in the isocortex (ISO) (SV40nls: 5526 ± 1156, ArgiNLS: 1798 ± 277) and olfactory area (OLF) (SV40nls: 1162 ± 211, ArgiNLS: 3753 ± 609), suggesting these two brain areas are major drivers of the differences in whole-brain cell counts.

To identify subregional differences in labeled cell densities, we used 1216 anatomical annotations provided by the Unified atlas^63^. First, we asked if the general distribution of labeled cell patterning across subregions differ amongst genetic tags (**Fig. 5e**). Linear regression analysis revealed a statistically significant positive correlation of ArgiNLS versus SV40nls labeled cell densities of subregions, suggesting an equivalent anatomical distribution of labeled cells across subregions. However, with the exception of 30 subregions, 98% contained greater cell densities when labeled with ArgiNLS (**Fig. 5e-f**).

Second, we screened for the identities of subregions significantly enriched in ArgiNLS-labeled cell density (**Fig. 5f bottom; 5g; Fig. S6-7; Dataset S1**). The most enriched gross regions – the ISO and OLF - contained a compilation of 60 (208/345; e.g., primary somatosensory area (S1)) and 23% (16/69; e.g., anterior olfactory area (AO), piriform cortex (Pir)) significant subregions, respectively (**Fig. 5g**; Supplemental Data Fig. 6). We found that ISO cortical enrichment was predominant in layers 2/3 to 5, with heavy concentration in areas related to sensory processing (**Fig. S7**). We additionally found significant enrichment of ArgiNLS-labeled cell density in subregions broadly located in every major division of the brain (HPF: 13/82 = 16%, e.g., dorsolateral entorhinal cortex (DLEnt), subiculum (SUB); CTXsp: 1/23 = 4%, e.g., intermediate endopiriform claustrum (IEn); CNU: 4/120 = 3%, olfactory tubercle (Tu); CB: 1/65 = 1.5%, e.g., lateral cerebellar nucleus parvicellular part (LatPC); TH: 6/94 = 6%, e.g., medial geniculate nucleus (MG); HY: 1/104 = 1%, medial mammillary nucleus (MM); MB: 8/109 = 7%, e.g., inferior colliculus (IC); and HB: 2/198% = 2%, e.g., Kolliker-fuse nucleus (KF)) (**Fig. 5g; Fig. S7**). We additionally performed an orthogonal, voxel-by-voxel analysis of differences in brain-wide cell density. Like our region-by-region analysis, significantly greater voxels were visualized across all major brain areas confirming wide-spread enrichment of labeled cell density by ArgiNLS (**Fig. 5h; Movie S1**).

### ArgiNLS-optimized applications in volumetric brain cell counting

Spatially integrated features of brain structure and function can be studied through volumetric imaging paired with genetically-encoded fluorescent labels. For example, many transgenic lines are available for, among other purposes, guiding FP labeling amongst a constellation of cell-types of choice^64^. Further, AAV technology is advancing rapidly as seen by recent serotype development directing FP labeling in retrograde, transsynaptic, or systemic fashions^65^. Since ArgiNLS optimizes single-cell FP labeling and segmentation, we established its utility for cell counting applications when incorporated into a variety of molecular genetic tools.

We first compiled an AAV vector toolkit consisting of multiple ArgiNLS-tagged FPs, including bright green-(AausFP1^66^ and mGreenLantern^67^), yellow-(mVenus ME^68^) and red-emitting (oScarlet^53^) variants within constitutive and conditional formats to offer experimental range (**Table S2**). We applied aqueous-based SHIELD clearing^69^ and axial swept LSFM imaging^70^ to demonstrate ease of ArgiNLS-assisted segmentation of single-cells and their total count quantification using 3D ML from native fluorescence in 3D volumes. We demonstrated ArgiNLS-assisted quantification by generating and injecting a retrograde AAV-EF1a-ArgiNLS-mKate2 virus into the basomedial amygdala (BMA) – an amygdalar structure involved in anxiety and fear modulation^71^ - to create an anatomical input map (**Fig. 6a-c**). Co-injected EGFP-expressing virus confirmed BMA injection placement, and native retro-ArgNLS-mKate2+ single-cells were readily identifiable at local and long-range distances from the injection site (**Fig. 6a**). 3D ML classification of brain-wide ArgiNLS-mKate2+ cells quantified at total of 2.4e^5^ BMA inputs that were mainly distributed continuously throughout the ipsilateral cortical subplate including amongst multiple amygdalar subregions (Amy) and claustrum (Cl), as well as within the subiculum of the hippocampus (**Fig. 6b**). Major long-range BMA input cells resided in 5 nodes including (from anterior to posterior) the accessory olfactory bulb (AOB), infralimbic cortex (IL), anterior paraventricular nucleus (aPVT) of the thalamus, contralateral amygdala, and ventral premammillary nucleus of the hypothalamus (PMv). The aPVT represented one of the largest long-range and singular thalamic node containing 5.6e^3^ input cells, highlighting the existence of a putative aPVT-BMA circuit motif of unknown behavioral function.

**Figure 6.**
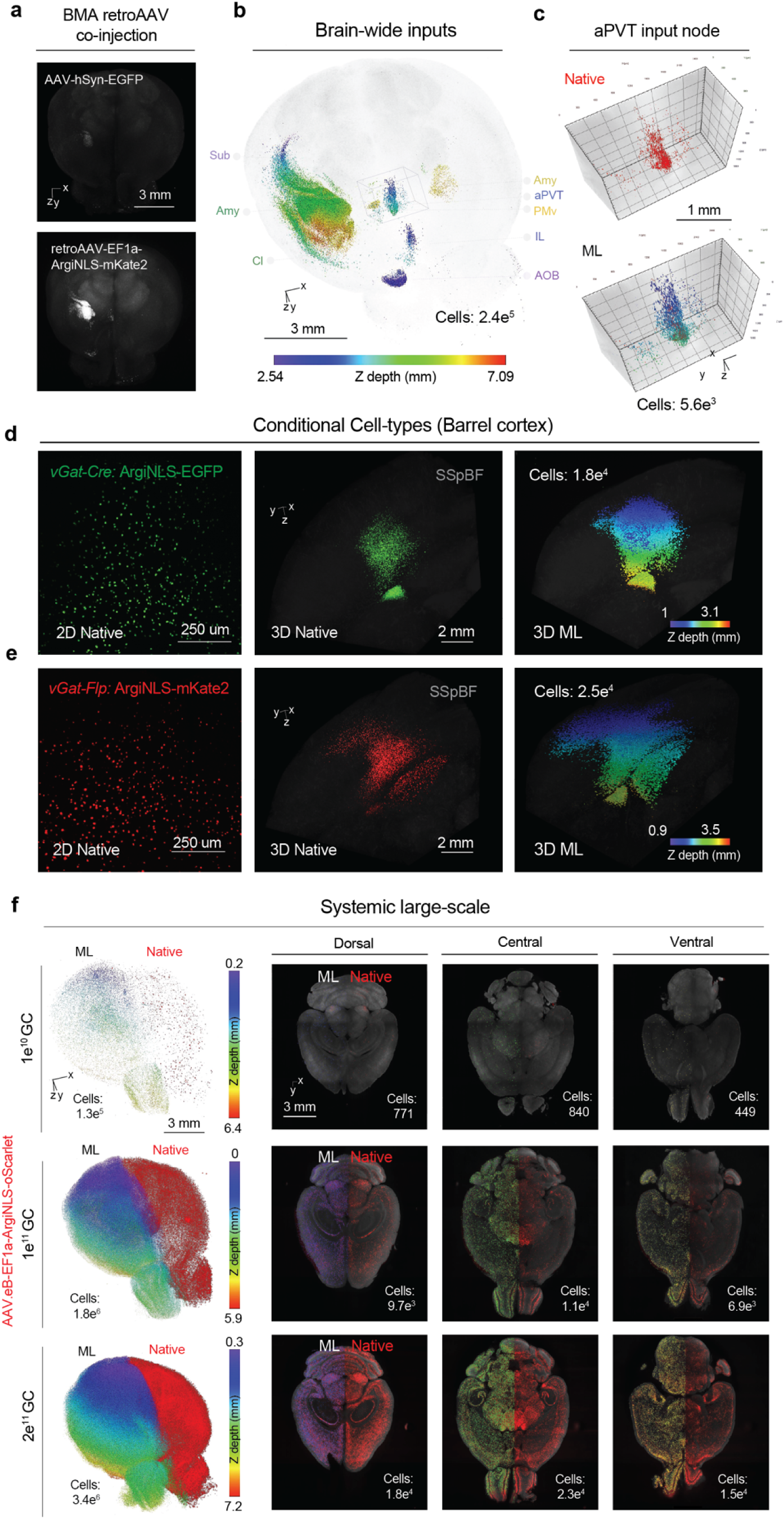
Optimized volumetric brain cell counting applications with ArgiNLS viral vectors. **a-c,** Example circuit mapping with ArgiNLS-mKate2-expressing retroAAV into the basomedia amygdala (BMA). **a,** Top: Whole-brain volumetric rendering of AAV-hSyn-EGFP native expression labeling the injection site in green channel. Bottom: Corresponding red channel image displaying native fluorescence of ArgiNLS-mKate2+ single-cell inputs. **b,** Volumetric rendering of single-cell brain-wide input segmentations and their total counts using 3D ML, gradient color coded by Z depth. Total BMA input cells are listed in the bottom right. Main anatomical areas of local and long-range inputs are listed to the left and right, respectively. Sub = subiculum; Amy = amygdala; Cl = claustrum; aPVT = anterior paraventricular nucleus of the thalamus; PMv = ventral premammillary nucleus; IL = infralimbic cortex; AOB = accessory olfactory bulb **c,** Single-cells (native fluorescence: top; ML segmentation: bottom) and their total count from the anterior PVT (aPVT) input node cropped and expanded from the outlined inlay in **b**. **d-e,** Conditional labeling of GABAergic single-cells in the barrel cortex using local (**d**) Cre- or (**e**) Flp-dependent AAV injections in *vGAT-Cre* or *vGAT-Flp* driver mice, respectively. Left: 2D confocal image stacks displaying native conditional single-cell labeling from separate example brain samples. Middle: Example volumetric renderings of native conditional single-cell labeling from separate brain samples. Right: Volumetric renderings of 3D ML-segmented single-cells and their total counts for each sample. **f,** Large-scale labeling of single-cells using systemic delivery of AAV.eB-Ef1a-ArgiNLS-oScarlet virus at 1e^10^, 1e^11^, or 2e^11^ GC payloads. Left: Whole-brain volumetric renderings of combined hemispheric native ArgiNLS-oScarlet fluorescence (left hemisphere) and 3D ML segmentation (right hemisphere) of single-cells from separate example brain samples per viral condition. Total hemispheric cell counts are listed in the bottom left. Right: 2D horizontal planes of each sample across (top to bottom) dorsal, central, and ventral positions. Images display native ArgiNLS-oScarlet fluorescence with overlaid 3D ML classification on the right hemisphere and total cell counts per plane listed in the bottom right.

Second, we generated and locally injected AAV.eB-CAG-FLEX-ArgiNLS-EGFP and AAV.eB-CAG-fDIO-ArgiNLS-mKate2 virus into the barrel cortex of *vGat-Cre* or *vGat-Flp* driver mice, respectively, to conditionally label and 3D segment GABAergic single-cells (**Fig. 6d-e**). High SNR-labeled single-cells were identically observed in Cre- or Flp-dependent conditions in 2D (**Fig. 6d-e, left**). In 3D, a clear distribution of single-cell-labeled viral spread was observed and devoid of extranuclear signal artifact in both Cre- and Flp-dependent conditions (**Fig. 6d-e, middle**), with 3D ML segmenting 1.8e^4^ and 2.5e^4^ GABAergic cells, respectively (**Fig. 6d-e, right**).

Finally, we generated and systemically delivered an AAV.eB-EF1a-ArgiNLS-oScarlet virus to demonstrate robust native fluorescence labeling of single-cells and their ease of segmentation of large cell amounts ubiquitously throughout the brain (**Fig. 6f; Movie S2**). To gauge sensitivity of detection levels enabled by ArgiNLS, we systemically delivered 3 amounts – 1e^10^ (1x), 1e^11^ (10x), and 2e^11^ (20x) - of viral GC payloads. 3D renderings of native signal displayed sharply contrasted single-cells distributed widely throughout the brain, and scaled with viral GC amount (**Fig. 6f, left**). 3D ML quantification of single hemispheres segmented 1.3e^5^, 1.8e^6^ (14-fold), and 3.4e^6^ (26-fold) single-cells for 1e^10^, 1e^11^, and 2e^11^ GC payloads, respectively, or 1 cell labeled for every 58,173 viral GCs delivered (**Fig. 6f, left; Supplemental Data Fig. 8a, left**). Expressing fold change values over the lowest 1e^10^ GC values indicated the utility of ArgiNLS-oScarlet to precisely label and segment 1.3-fold cell count increases for every 1-fold increase in viral GC delivered, within the 1 to 20-fold (i.e.,1e^10^-2e^11^) GC range tested (**Fig. S8a, right**). This total single-cell detection by viral GC relationship was maintained evenly throughout the brain as observed in cell counts across separate horizontal planes located in dorsal, central, and ventral anatomical positions (**Fig. 6f, right; Fig. S8b**).

## Discussion

Volumetric imaging methods can be used to unveil spatially integrated representations – or maps – of brain structure and function at the direct level of single-cells. Pivotal to this approach is the ability of genetically-encoded fluorescent labels to discriminate single-cells within dense and complex brain tissue. Here, we developed and characterized a new nuclear-localizing genetic tag strategy named ArgiNLS that optimizes FP labeling of single-cells, without off-target effects on cellular physiology or animal behavior. By extension, ArgiNLS provides seamless compatibility and enrichment of ML-automated brain cell segmentation across small-to-large scale image datasets of various applications, demonstrating its use as an improved single-cell resolution brain mapping genetic tool.

Unlike other recently developed synthetic NLS approaches created for enhanced genome-editing^37,38,72^, the ArgiNLS approach for image segmentation is based on a well-understood molecular foundation. This approach uses a poly-R-based rerouting mechanism to reduce extra-nuclear NLS-tagged FP localization by attracting it into the nucleolus. Optimal tag efficacy of ArgiNLS occurs when the BP cNLS sequence is combined with a specific poly-R length (R7) that titrates FP localization into a balanced nucleoplasmic/nucleolar composition with optimal native brightness. MP cNLS combinations fail to reduce extra-nuclear artifact in combinations up to 11 Rs, suggesting that the function of R7 synergism on ectopically-fused NLS activity requires sequence and/or length characteristics of a BP NLS. The function of this configuration within cells outside of the CNS and amongst varied species was not determined in our current study, and therefore context-specific modification may be required. However, as demonstrated in mouse brain here, the ArgiNLS strategy lays a robust framework for tailoring function through the parameterization of BP NLS sequence selection and/or poly-R lengths.

The nature of the poly-R7 component of ArgiNLS additionally provides functional fidelity across a wide variety of FPs, that can provide spectral flexibility for single-cell labeling, and potential nuclear localization of other non-FP proteins, through its dual role as a protein linker. This capability was demonstrated on an initial group of spectrally separate FPs. However, further validation amongst a larger FP selection and with non-FP proteins for applications outside of image segmentation is necessary to strengthen the generalization of this linker-like modularity. Additionally, while our current ArgiNLS strategy applies N terminus tags to FPs, we did not test linker-like capability or function when attached to the C terminus, thereby limiting its current principles of use to a single fusion location.

NLS performance of ArgiNLS *in vivo* demonstrates several advantages over current SV40nls tag or H2B fusion single-cell labeling approaches. First, ArgiNLS causes FP nuclear restriction that is equally robust whether an AAV is delivered locally or systemically to the brain. ArgiNLS eliminates background fluorescence-producing, extra-nuclear signal that is consistently observed with SV40nls. ArgiNLS elicits consistent nuclear-restricted FP labelling across major cortical cell classes. This property enables improved ML image classification of intact whole brain tissue, which is characterized by a reduction in model training requirements and coupled to increased precision and recall of segmentations. Unlike H2B fusion^35^, these benefits were not associated with deleterious effects to neural physiology or animal behavior. Therefore, ArgiNLS is appropriate for genetic labeling experiments of single-cells that require concomitant behavioral procedures paired with AAV infection. Future time-course studies of *in vivo* expression durations longer than studied here (i.e., 3-4 weeks), will help understand the total range of temporal compatibility with cellular and animal health.

The in vivo use of ArgiNLS highlighted how the selection of genetic labelling strategy can significantly influence whole-brain single-cell segmentations, with an effective doubling of cell counts across subregions located in all major brain structures. The exact nature of this improvement is unclear; however, it is possibly related to depletion of extra-nuclear FP localization that enhances single-cell SNR. Additionally, since a fraction of wild-type FP molecules misfold or do not form a proper chromophore^73,74^, poly-R driven interaction with nucleolar ribosomal RNA^46^ and/or nucleolus-dependent changes of pH could provide a stabilizing influence in aggregation or folding of ArgiNLS-tagged FPs, leading to more single-cells detected.

Finally, ArgiNLS can be applied in various volumetric cell labeling applications to resolve cell counts seamlessly and rapidly with 3D ML segmentation. Our proof-of-concept experiments begin to demonstrate ArgiNLS compatibility with, and enhancement of, anatomical input mapping, conditional segmentation of cell-types, and mapping a wide range of titrated cell counts within 3D brain volumes using various AAV vectors. However, ArgiNLS may also be incorporated into other molecular genetic tools that will provide improvements to single-cell counting applications including intersectional mapping^74^ with or without multiplexing of spectrally separate ArgiNLS-tagged FPs, transsynaptic anatomical tracing^65^, activity-dependent reporting^75^, and transgenic reporter mouse lines for genomically-encoded expression^64^. Indeed, while we demonstrate improved single-cell labeling capacity of ArgiNLS over other tags, our *in vivo* characterization was solely based off AAV expression. The creation of future ArgiNLS-tagged expression vectors for gene delivery from other commonly used viral vectors^76^, as well as gene-targeting vectors for the creation of reporter mouse lines, will help expand optimized single-cell labeling and segmentation amongst broad genetic delivery options.

In conclusion, the simple yet powerful ArgiNLS genetic tag strategy described here advances streamlined quantification of single-cells from small-to-large scale brain image datasets with enriched labeling capacity and enhanced ML capabilities.

## Materials and Methods

### Mice

Adult male and female mice (8-12 weeks old) were used for all experiments unless otherwise noted. Animals were group housed on a reverse 12-hr light-dark cycle (0700 OFF/1900 ON) and had access to water and food ad libitum. All experimental procedures were performed in accordance with University of Washington IACUC guidelines. Wild-type and transgenic mice lines were purchased from the following vendors: C57Bl6/J (JAX, 000664); CD-1 (Charles River, Crl:CD1(ICR)); vGlut1-Cre (Slc17a7-IRES2-Cre-D; JAX, 023527); vGat-Cre (Vgat-ires-cre knock-in (C57BL/6J); JAX, 028862); vGat-Flp (Slc32a1-IRES2-FlpO-D knock-in; JAX, 031331). Descriptions of stereotaxic and retroorbital viral injections, electrophysiology, and behavioral testing are provided in **SI Methods**.

### Fluorescent protein-expressing AAV Vector construction

Untagged and NLS-tagged fluorescent proteins were cloned through PCR amplification using standard primers with or without extension overhangs (Integrated DNA Technologies). The monopartite SV40nls coding tag sequence used was from amino acids 126-132 of large T antigen (Betapolyomavirus macacae; NCBI Reference sequence: NP_043127.1). Amino acids 156-172 of the bipartite mouse nucleoplasmin2 NLS (NCBI accession number: NP_001398932.1) was used for NPM2nls. Isolectric point (pI) and net chrarge (z) calculations of each poly-R7 modified or unmodified cNLS protein tag was determined through the open source Prot pi online tool (http://protpi.ch). Starting Addgene plasmids and cloning strategies for construction are described in **SI Methods.**

### Characterization of nuclear localization in vitro

Immortalized embryonic mouse hippocampal cells (Cedarlane; mHippoE-1; cat. CLU198)) or mouse Neuroblastoma 2A cells (ATCC; CCL-131) were cultured in DMEM (high glucose and L-glutamine; Gibco 11965–092), supplemented with 10% fetal bovine serum (Gibco; 1600) and 1% penicillin-streptomycin (Invitrogen; 15140122) at 37°C in 5% CO_2_. Cells were sub-cultured following manufacturer’s instructions. For transient DNA transfections, cells were seeded onto Poly-L-lysine-coated glass coverslips (Electron Microscopy Science, 72292-08) in a 24-well plate format. Once cells reached ~80% confluence, transient transfections were performed in duplicate with 250 ng of plasmid DNA per construct using Lipofectamine 3000 transfection reagent (ThermoFisher Scientific, L3000015). After 48 h, cells were fixed with formalin, counterstained with DAPI, and mounted onto glass slides. All NLS tagged variants were co-transfected with a spectrally separate untagged fluorescent protein to fluorescently label total cellular area in the following combinations: all NLS tagged EGFP constructs with AAV-EF1a-mKate2; all NLS tagged RFP constructs with p-EGFP-N1 (Addgene; 6085-1). To qualitatively verify Cre-dependent ArgiNLS variant expression, each plasmid was co-transfected with pCAG-Cre (Addgene; 13775), respectively (data not shown). To qualitatively verify Flp-dependent ArgiNLS-mKate2 expression, we PCR-amplified and subcloned FlpO from PGK-NLS-FlpO-bpA (a gift from Dr. Phillipe Soriano^77^) into ptCAV-12a^78^ at AgeI/EcoRV restriction sites to create the FlpO-expressing ptCAV-FlpO-12a which was used for co-transfections (data not shown). Images of all transfection conditions were acquired with a Keyence BZ-X800E inverted fluorescence microscope using a 20x/0.75 NA objective (Nikon). The effects of experimental NLS and ArgiNLS tags on nuclear localization of fluorescence was quantified amongst single-cells (n=5 randomly selected cells/3 replicate cultures or otherwise noted) in a blinded fashion using ImageJ. Accordingly, manual segmentations of the nucleus as defined by DAPI fluorescence, and the whole cell as defined by untagged co-transfected plasmid fluorescence, were created in synchronized duplicate images of the target channel. Corrected total nuclear or whole cell fluorescence (CTNF or CTCF, respectively) were separately calculated using area, mean, and integrated density measurements within the following equation: CTF = Integrated density - (Area * Mean background). Percent nuclear localized fluorescence was then calculated for each cell as CTNF/CTCF * 100.

### Adeno-associated virus production

#### Php.eB and retroAAV2 packaging plasmid construction

A single packaging plasmid to produce PHP.eB viruses, pDG_PHP.eB, was constructed by subcloning the PHP.eB capsid from pUCmini-iCAP-PHP.eB (Addgene, 103005) into pDG1 plasmid to replace the AAV1 capsid at ClaI/SwaI restriction enzyme sites. The final vector was produced at 1mg/mL by Nature Technologies (Lincoln, Nebraska). A single packaging plasmid for retroAAV2 was created by subcloning the retroAAV2 capsid^80^ into the pDG plasmid in a manner identical to pDG_PHP.eB to create pDG_retroAAV2. The pDG_PHP.eB plasmid is publicly available through Addgene. AAV generation and genome copy (GC) titration methods are provided in **SI Methods**.

### Characterization of ArgiNLS-EGFP nuclear localization in vivo

Local AAV-injected mice were terminally anesthetized with isofluorane and transcardially perfused with phosphate-buffered saline (PBS) followed by 10% neutral buffered formalin (Sigma, HT501128). Brains were extracted, post-fixed overnight in 4 C, and cryoprotected in 30% sucrose at 4 C. Cryoprotected brains were histologically sectioned in coronal orientation on a freezing cryostat (Leica Biosystems) at 50 um in 6 equal series. Sections were counterstained with DAPI (Millipore Sigma; 10236276001) and then mounted with VectaShield mounting media (Vector Laboratories; H-1400-10). Mounted sections were imaged using a Ziess LSM880 confocal microscope. Confocal images were captured with either 10x/0.45 NA (Zeiss) or 40x/1.2 NA (Zeiss) objectives. All scans were acquired in 16-bit with 2.05 ms pixel dwell time and line 8 averaging in a 1024 x 1024 dimension. 3-channel DAPI/EGFP/tdTomato images were captured using the following excitation/emission filters: DAPI: 410-585 nm; EGFP: 493-564 nm; tdTomato: 566-691 nm. Consistent laser power and gain settings were used across all conditions. 40x scans used for quantification were acquired in Z-stacks consisting of 3 um optical steps across 15 um total tissue depth. Maximal z-projections were created in ImageJ and used for quantification. The effects of SV40nls and ArgiNLS tags on cellular localization of EGFP fluorescence was quantified in EGFP+/tdTomato+ single cells (n=5 randomly selected cells/3 vGlut1-Cre or vGat-cre animals) from 40x images in a blinded fashion using ImageJ in a manner similarly described for *in-vitro* quantification. Briefly, manual segmentations of the nucleus as defined by DAPI fluorescence, and the soma as defined by Cre-dependent tdTomato fluorescence, were created in synchronized duplicate images of the target EGFP channel. Since we observed different levels of extracellular background fluorescence at the injection site amongst SV40nls- and ArgiNLS-EGFP injections, we used DAPI-guided extracellular background fluorescence images and measurements from the uninjected contralateral side of the SSpBF for background subtraction. These values were also used to compare against the extracellular fluorescence background measurements taken within each injection site of all animals to perform ipsilateral versus contralateral extracellular background fluorescence comparisons. Axonal fluorescence was quantified from additional 20x image stack z maximal z-projections acquired in the corpus callosum from *vGlut1-Cre* samples. For quantification, we applied an adaptive manual threshold to isolate measurements to axons only using the Huang method in ImageJ on 608 x 1024 FOVs. Corrected total axonal fluorescence (CTAF) was then calculated for each channel as Integrated density – (Area * mean background). EGFP+ CTAF was internally normalized for each sample to *vGlut1-Cre*:tdTomato+ CTAF and expressed as a percent.

### ML classification performance in 2D images

Mice RO-injected with AAV.eB-EF1a-ArgiNLS-EGFP or AAV.eB-EF1a-Sv40nls-EGFP were terminally anesthetized with isofluorane and transcardially perfused and histologically processed as described above. Single-plane epifluorescent images of entire coronal planes at AP 2.11 and −0.52 were acquired in montages with a Keyence BZ-X800E inverted fluorescence microscope using a 10x/0.45 NA objective (Nikon). Consistent DAPI and EGFP channel exposure settings were applied across conditions. Montages were stitched using BZ-X800 Analyzer software (Keyence, v 1.1.1.8) and composite images were created in ImageJ and exported as TIFF images. Machine learning classifiers for ArgiNLS- and SV40nls-EGFP labels were created using random forest classification with QuPath^81^ software applied to composite TIF images (see **SI Methods**).

### 3D volumetric characterization of ArgiNLS-EGFP performance

Two independent approaches were applied to systematically quantify and characterize single-cell labeling across the intact mouse brain with ArgiNLS-tagged FPs: 1) iDISCO+ solvent-based brain clearing and EGFP staining based on methods previously described^9,13^ with modification, and 2) SHIELD aqueous-based brain clearing (see **SI Methods**).

### Whole-brain image processing

#### Brain-wide SV40nls versus ArgiNLS-tagged EGFP cell count comparisons

Volumetric LSFM-acquired images were processed for whole-brain template registration, automated cell segmentation, and regional segmentation as previously described^9^ with modifications (see **SI Methods**).

### 3D Visualization and ML

Zeiss arivis Pro (formerly, Vision4D) software was used to generate 3D renderings of brain samples throughout the manuscript (see **SI Methods**).

## Author Contributions

E.R.S/S.A.G conceived of the project and E.R.S developed the genetic tag strategy. E.R.S performed all stereotaxic injections and confocal microscopy. E.R.S/L.K performed *in vitro* experiments and molecular cloning. Y.Z/B.B performed histology. E.R.S/J.N performed retro-orbital injections for all experiments. E.R.S/J.N performed epifluorescent imaging. J.N implemented QuPath for 2D cell segmentation testing. J.N/Y.Z/L.K performed manual annotations and F1 scoring for 2D cell segmentation. A.D.M performed iDISCO+ brain clearing and staining and LSFM imaging. E.R.S/A.D.M implemented ClearMap volumetric image processing pipeline. M.C.M/M.B performed behavioral battery experiments. B.J/K.S.G performed electrophysiology recordings and analysis. E.R.S performed all analyses. E.R.S/S.A.G wrote the manuscript.

## Acknowledgments

We would like to acknowledge support from the University of Washington Center of Excellence in Addiction Pain and Emotion (NAPE) Molecular Genetics Resource Core (P30DA048736). We would like to thank Valerie Tsai, Aida Chan, Axel Salazar, and Kevin Bai for their technical support. We thank the entire Golden and Heshmati Laboratories for their discussions and critiques of the manuscript.

