## Supplemental Information for "An arginine-rich nuclear localization signal (ArgiNLS) strategy for streamlined image segmentation of single-cells"

**This PDF file includes:**

Supporting text

Supporting methods

Figures S1 to S8

Tables S1 to S2

Legends for Movies S1 to S2

Legends for Datasets S1

SI References

**Other supporting materials for this manuscript include the following:**

Movies S1 to S2

Datasets S1

Supporting Information Text

**Page 7:** ML segmentation revealed a higher density of cell counts using ArgiNLS-EGFP (**Fig. 5b; Fig. S5c)**, with both classifiers similarly segmenting raw EGFP-stained cells (**Supplemental Data Fig. 5d**). We quantified the performance of each 3D classifier using systematic F1 comparisons of ML segmentations versus human annotations (i.e., ground truth) in matching counting box FOVs located amongst 5 separate brain regions of varying cell density, including the frontal cortex (FC), primary motor cortex (MO), thalamus (TH), hippocampus (HIP), and hindbrain (HB) (**Fig. S5e**). As anticipated due to training optimization (**Fig. S5a-b**), no significant differences in F1 scores – or classifier performance - were found across counting box brain areas, with final composite F-scores reaching 0.85 for SV40nls, and 0.83 for ArgiNLS (**Fig. S5e**). We then quantified differences in cell density by comparing mean ground truth and ML classifier cell counts as a function of genetic tag. ArgiNLS infected brains contained a significantly greater number of human-counted stained cells across all counting boxes, delivering a significant composite fold increase of 2.8 (**Fig. S5f**). Similarly, ArgiNLS infected brains contained a greater number of ML-segmented cells across all FOVs that was significant for FC and MO only, providing a significant composite fold increase of 3.4 (**Fig. S5g**).

**Supporting Information Methods**

*Stereotaxic viral injections*

All stereotaxic surgeries were performed under aseptic conditions. Briefly, animals were anesthetized with isoflurane prior to and during placement and surgery in a stereotaxic instrument (David Kopf Instruments). Scalp incisions were made to expose the skull surface and virus was injected intracranially using a 32 gauge 0.5 ul Neuros syringe (Hamilton Co.) at the following coordinates from bregma of the mouse brain: SSpBF: -1.67 anterioposterior, 2.75 mediolateral, -0.9 dorsoventral; BMA: 1.55 anterioposterior, 2.7 mediolateral, -5.2 dorsoventral. For NLS nuclear fluorescence comparisons in defined cell-types, 400 nl of AAV.eB-EF1a-ArgiNLS-EGFP (7x10^12^ GC/ml) or AAV.eB-EF1a-SV40nls-EGFP (7x10^12^ GC/ml) was co-injected with 100 nl of AAV.eB-CAG-FLEX-tdTomato (Addgene, 28306; 7x10^13^ GC/ml) into the SSpBF of homozygous vGlut1-Cre or vGat-Cre mice at a 0.1 ul/min delivery rate. For qualitative characterization of conditional ArgiNLS-tagged fluorescent proteins, 500 nl of AAV.eB-CAG-FLEX-ArgiNLS-EGFP (1x10^12^ GC/ml) and AAV.eB-CAG-fDIO-ArgiNLS-mKate2 (1x10^12^ GC/ml) virus was injected into the SSpBF of separate heterozygous vGat-Cre or vGat-Flp mice at a 0.1 ul/min delivery rate. For characterization of retrograde ArgiNLS-tagged fluorescent protein expression, 200 nl of retroAAV2-EF1a-ArgiNLS-mKate2 (1.5x10^13^ GC/ml) virus was injected into the BMA of C57Bl6/J mice at a 0.02 ul/min delivery rate. All surgerized animals received 1 mg/ml ketoprofen analgesic and 500 ul saline subcutaneously followed by recovery in their homecage placed above a warm heating pad. All animals receiving stereotaxic injections were sacrificed for histological analysis at 2 weeks post-injection.

*Retroorbital viral injections*

Unless otherwise noted, 1x10^11^ GC of AAV.eB virus was diluted up to 100 ul with sterile saline and injected intravenously into the retro-orbital (RO) sinus of mice receiving brief isofluorane anesthesia. All RO-injected animals were sacrificed at 3 weeks post-injection for histological analysis.

*Fluorescent protein-expressing AAV Vector construction*

The following Addgene plasmids were used for PCR templates in AAV vector construction: pAAV-CAG-NLS-GFP (104061) for all EGFP variants; pAAV2-CAG-3xNLS-AausFP1 (191096); pcDNA3.1-mGreenLantern (161912); pEB1-mVenus ME (103989); pAAV-EF1a-oScarlet (137135); pAAV-RAM-d2TTA::TRE-NLS-mKate2-WPREpA (84474) ; pmiRFP670-N1 (79987); and pCS-H2B-EGFP (53744) . For constitutive expression AAV construction, fluorescent proteins and variants were subcloned into pAAV-Ef1a-DIO-Synaptophysin-GCaMP6s (105715) at XhoI/EcoRV restriction enzyme sites. For Cre-dependent expression AAV construction, all fluorescent proteins and ArgiNLS-tagged variants were subcloned into pAAV-CAG-FLEX-GFP (59331) at KpnI/XhoI restriction enzyme sites. For Flp-dependent expression AAV construction, ArgiNLS-mKate2 was subcloned into pAAV-CAG-fDIO-mNeonGreen (99133) at KpnI/XhoI restriction enzyme sites. ArgiNLS-GCaMP6s and ArgiNLS-EGFP variants containing less or more than 7 R’s were gene synthesized and cloned into the pAAV-Ef1a-DIO-Synaptophysin-GCaMP6s vector at Genscript (Piscataway, NJ). All cloned insert sequences of final plasmids used in the manuscript were sequence verified by Sanger sequencing (Genewiz, Seattle, WA). Full AAV plasmid sequences and maps are made publicly available through Addgene.

*AAV generation*

HEK293T cells were transfected with 25 μg AAV vector plasmid and 50 μg packaging vector (pDG_PhP.eB or pDG2_retroAAV2) per 15 cm plate. Two days after transfection, cells were harvested and subjected to three freeze–thaw cycles. The supernatant was transferred to a Beckman tube containing a 40% sucrose cushion and spun at 27,000 rpm overnight at 4°C. Pellets were resuspended in CsCl at a density of 1.37 g/ml and spun at 65000 rpm 4 hours at 4°C. 1 ml CsCl fractions were run on an agarose gel, and genome-containing fractions were selected and spun at 50000 rpm overnight at 4°C. The 1 ml fractions were collected again, and genome containing fractions were dialyzed overnight. The filtered solution was transferred to a Beckman tube containing a 40% sucrose cushion and spun at 27,000 rpm overnight at 4°C. The pellet (containing purified AAV) was resuspended in 100 μl 1× HBSS. Virus was aliquoted and stored at -80 ° C until use.

*AAV genome copy (GC) titration*

Two methods were employed to titrate viral DNA genome copies of AAV preparations: dPCR by Stanford’s Gene Vector and Virus Core for initial constitutive and conditional viral preparations, and in-house SYBR green-based qRT-PCR for subsequent constitutive preparations. For dPCR, the Qiacuity 5-plex system was employed which uses random distribution of template in available partitions to detect and measure fluorescence emission from target amplicons. The virus is was first made DNA ready by adding 8ul of Biosearch Quickextract solution to 2 ul of sample and thermocycling at 65C for 6min and 98c for 2min. Extracted DNA was then combined at 100x, 10,000x, and 5,000,000x dilutions with Qiacuity ProbePCR mastermix and XFP primers (forward: agcaaagaccccaacgagaa; reverse: ggcggcggtcacgaact), and reaction mixes were subjected to a dPCR thermocycling program: 95 C for 2 min, and 50 cycles of 15 sec at 95 C and 30 sec of 60 C. Earlier determined titer of AAV-45 was included as internal control along with a No Template Control.  The concentration 'copies/ul' is then used to calculate the genome copies per mL. For qRT-PCR, 5 ul of AAV at multiple dilutions (5000x, 25000x, 125,000x, 625,000x, and 3,125,000x) or linearized standard plasmid DNA (1e^4^-1e^8^ molecules) were combined into master mixes containing SsoAdvanced Universal SYBR Green (Bio-Rad, 1725271) and primer sets targeting either the WPRE3 sequence (forward: CTGGTTAGTTCTTGCCACGG; reverse: AATTGTCAGTGCCCAACAGC) for titration of constitutive viruses. 20 ul reaction mixes were then subjected to the following thermal cycling protocol using a Bio-Rad CFX Connect Real-Time PCR System: 2 min at 98 C, and 38 cycles of 15 sec at 98 C and 20 sec at 60 C. Viral genome copies/ul were calculated in Microsoft Excel based off of plasmid DNA standard curves that passed quality control of an R^2^ of ~.99 and PCR efficiencies of ~100%.

*QuPath Classifier Training*

We used 25 px x 25 px (625 px^2^) tile FOVs labeled as ‘DAPI’ or ‘GFP’ as training input data. The object classification function was used to create classifiers with the following train object classifier parameters: object filter = all detections; classifier = random trees (RTrees), features = all measurements, classes = all classes, and training = unlocked annotations. Cell detection parameters consisted of: Setup parameters detection channel = DAPI; nucleus parameters background radius = 0 px; median filter radius = 0 px; sigma = 3 px; minimum area = 10 px^2^; maximum area = 200 px^2^; intensity threshold = 50 and split by shape; cell expansion = 0 px; general parameters = smooth boundaries and make measurements. The live update function was used to observe classifier training, and each training iteration included the addition of 2 training tiles. The first training iteration for each classifier included both a ‘DAPI’ and ‘GFP’-labeled training ROI.  Total iterations to reach maximal performance for the ArgiNLS classifier was 10 and included a combination of 7 DAPI and 2 GFP training ROIs. Total iterations to reach maximal performance for the SV40nls classifier was 28 and included a combination of 22 DAPI and 6 GFP training ROIs.

*Classifier Performance Quantification*

Additional 100 px x 100 px (10,000 px^2^) FOVs were created (4 per AP) to quantify classifier performance at every training iteration. FOV placement varied ranging from regions with small to large density of labeled cells. Labeled cells in each FOV were manually segmented separately by 3 expert raters, and classifier training was repeated, as described above. Both manual and automated segmentations were overlaid for each FOV in QuPath and manual only segmentations, classifier only segmentations, and co-segmentations were hand scored for each FOV at every training iteration. Data was recorded in Microsoft Excel and used to calculate precision, recall, and F1 scores.

*iDISCO+ whole-brain processing*

Mice RO-injected with AAV.eB-EF1a-ArgiNLS-EGFP (n=4) or AAV.eB-EF1a-Sv40nls-EGFP (n=4) were sacrificed and brains were post-fixed as described above. We performed iDISCO+ beginning with a pretreatment of 1x PBS washes (3 x 30 min; RT) followed by dehydration using ascending incubations of methanol (MeOH; Sigma, 34860-4L-R) concentrations (1 h each step; RT rotating): 20%, 40%, 60%, 80%, 100%, and 100%. Samples were next delipidated with a 66% dichloromethane (DCM; Sigma, 270997-12X100ML)/34% MeOH solution (ON, RT rotating), and washed the following day in 100% MeOH (2 x 1 h, RT rotating). Brains were then bleached in a chilled solution consisting of 5% hydrogen peroxide (30% Hydrogen peroxide solution; Sigma, 21673-500ML)/methanol (ON,4 C). Bleached brains were then rehydrated using descending incubations of MeOH concentrations (1 h each step; RT): 80%, 60%, 40%, 20%, and finally washed in 1x PBS buffer containing 0.5% TritonX-100 (Sigma, T9284-1L) (2 x 1 h; RT rotating). To immunostain brains for EGFP, we first permeabilized pre-treated samples in a 1x PBS buffer containing 0.5% TritonX-100/1.4% glycine (Sigma, G7126-1KG)/20% dimethyl sulfoxide (DMSO; Sigma, D128-1) (2 days, 37 C rotating) followed by blocking of non-specific binding through incubation with a 1x PBS buffer containing 0.5% TritonX-100/6% normal donkey serum (NDS; Jackson ImmunoResearch, 017-000-121)/10% DMSO (2 d, 37 C rotating). EGFP primary antibody (chicken polyclonal; Aves, GFP1010) labeling was next performed by applying a 1:200 dilution in a 1x PBS buffer containing 0.5% Tween-20/10ug/mL heparin (Sigma, H3393-250KU)/ 3% NDS/5% DMSO (7 days, 37 C rotating). Excess antibody was then washed with 1x PBS buffer containing 0.5% Tween-20/10ug/mL heparin (4-5 times x 2 days, 37 C rotating). Secondary antibody (donkey anti-chicken Alexa Fluor-647-conjugated AffiniPure F(ab’)2; Jackson ImmunoResearch; 703-606-155; Lot: 157044) staining was then performed by applying a 1:100 dilution in a 1x PBS buffer containing 0.5% Tween-20/10ug/mL Heparin/ 3% NDS/5% DMSO (7 days, 37 C rotating). Excess secondary antibody was washed with 1x PBS buffer containing 0.5% Tween-20/10ug/mL heparin (4-5 times x 3 days, 37 C rotating). Tissue clearing then began with dehydration using ascending incubations of MeOH concentrations (1 h each step; RT rotating): 20%, 40%, 60%, 80%, 100%, and 100% followed by delipidation with a 66% DCM/34% MeOH solution (3 h, RT rotating). Next, MeOH was washed with 100% DCM (2 x 15 min; RT rotating). Finally, samples were index matched for at least 1 day in DiBenzyl Ether (DBE; Sigma, 108014-1KG) prior to imaging.

*iDISCO+-processed volumetric imaging*

EGFP-stained and cleared whole-brains were imaged in DBE using an UltraMicroscope II light-sheet fluorescent microscope (LSFM) with Infinity Corrected Objective Lenses (Miltenyi Biotec). Our microscope configuration consisted of a 1.1x/0.1 NA MI PLAN objective (LaVision BioTec), non-corrected dipping cap, and an Andor Zyla sCMOS camera. All samples were secured on a 3D-printed sample platform and positioned in horizontal orientation with the cortical surface facing up. 2-channel image acquisition was performed in 1 FOV tile for autofluorescence (Excitation: 488 nm laser, Emission: 535/43 bandpass filter) and EGFP stain (Excitation: 647 nm laser, Emission: 690/50 bandpass filter) in separate scans (scan order: z-x-y) at a near-isotropic pixel resolution of 5.91 um X x 5.91 um Y x 6 um Z. We applied the following fixed acquisition parameters across all samples using ImspectorPro software (Miltenyi Biotec; v 7.1.16): laser power = 20 (488 channel), 30 (647 channel); exposure = 100 ms (488 channel), 136 ms (647 channel); sheet NA = 0.16; sheet thickness = 3.89 um; sheet width = 100%; zoom = 1x; dynamic horizontal focus = 5 (647 channel only); dynamic horizontal focus processing = contrast adaptive; merge light-sheet = blend.

*SHIELD whole-brain processing*

We followed the SmartClear full active pipeline protocol (LifeCanvas Technologies, v5.05) for aqueous-based brain clearing and mounting. Accordingly, RO- or locally-injected mice were sacrificed and brains were post-fixed as described above. Brain samples were first incubated in freshly prepared SHIELD OFF solution, consisting of SHIELD buffer and epoxy solution (4 days, 4 C shaking), followed by SHIELD ON buffer (1 day, 37 C shaking). Samples were next equilibrated in Delipidation buffer (ON, RT shaking). Prior to active clearing, the SmartClear II Pro electrophoresis system (LifeCanvas Technologies) containing Delipidation buffer was equilibrated to 42 C and new membranes were installed. Samples were then transferred into mesh bags and placed into clearing chambers. Active clearing was performed on 4-sample batches using the Gentle Mode program (1500 mA/42 C; 1 day). After clearing, samples were removed from mesh bags and index matched first in 50% EasyIndex (1 day; 37 C slow shaking) followed by 100% EasyIndex (1day; 37 C slow shaking). Index matched samples were embedded on custom 3D-printed sample mounts in EasyIndex containing 2% w/v agarose (Sigma, A5030), and incubated post-embedding in 100% EasyIndex (ON, 37 C). Embedded samples were stored in 100% EasyIndex (RT) in the dark until imaging.

*SHIELD-processed volumetric imaging of native fluorescence*

Native signal from aqueous-based cleared brains were imaged horizontally using the SmartSPIM LSFM (LifeCanvas Technologies)^70^ at 4 um isotropic pixel resolution. We imaged all samples in 488 and 563 channels to acquire native fluorescence from AAV expression from green- or red-emitting FPs, and background fluorescence in the channel without AAV expression for 3D rendering purposes. Laser power and acquisition settings were optimized individually towards each proof-of-concept sample, whereas settings for titrated AAV.eB-EF1a-ArgiNLS-oScarlet samples were held constant across conditions.

*Whole-brain image processing*

First, we employed the Unified brain template and atlas^63^ for registration and regional segmentation, respectively. Second, we applied ML classification of single-cells from SV40nls-EGFP or ArgiNLS-EGFP stains through the creation of separately trained classifiers using random forest classification algorithm through ilastik software^62^. Classifier Performance was quantified in FOVs in regions with small to large density of labeled cells as similarly described previously on non-ML segmented datasets^9,21^. Labeled cells in each FOV were manually segmented separately by 3 expert raters. Data from ImageJ processing was recorded in Microsoft Excel and used to calculate precision, recall, and F1 scores.

*Electrophysiology*

*vGlut1-Cre* mice injected with ArgiNLS-EGFP and Cre-dependent tdTomato virus were allowed 3-4 weeks recovery after surgery to allow for viral expression. All solutions were continuously bubbled with O_2_/CO_2_. Coronal (200 um) brain slices were prepared from 11-13 week old mice in a slush NMDG cutting solution (in mM: 92 NMDG, 2.5 KCl, 1.25 NaH_2_PO_4_, 30 NaHCO_3_, 20 HEPES, 25 glucose, 2 thiourea, 5 Na-ascorbate, 3 Na-pyruvate, 0.5 CaCl_2_, 10 MgSO_4_, pH 7.3–7.4^82^. Slices recovered for ~12 min in the same solution in 32°C water bath. Slices were then transferred to a room temperature HEPES-aCSF solution (in mM: 92 NaCl, 2.5 KCl, 1.25 NaH_2_PO_4_, 30 NaHCO_3_, 20 HEPES, 25 glucose, 2 thiourea, 5 Na-ascorbate, 3 Na-pyruvate, 2 CaCl_2_, 2 MgSO_4_). Slices recovered for an additional 60 min. Whole-cell patch-clamp recordings were made using an Axopatch 700B amplifier (Molecular Devices) with sampling at 10KHz and filtering at 1 kHz. EGFP+ and/or tdTomato+ cells were visually identified via fluorescence for patching with electrodes at 4–6 MΩ. Series resistance was monitored during all recordings with changes in resistance +10% qualifying the cell for exclusion. Recordings were made in aCSF (in mM: 126 NaCl, 2.5 KCl, 1.2 NaH_2_PO_4_, 1.2 MgCl_2_ 11 D-glucose, 18 NaHCO_3_, 2.4 CaCl_2_) at 32°C continually perfused over slices at a rate of ~2 ml/min. Barrel cortex glutamate neurons were identified by fluorescence. Electrodes were filled with an internal solution containing (in mM): 130 potassium gluconate, 10 HEPES, 5 NaCl, 1 EGTA), 5 Mg2+/ATP, 0.5 Na+-GTP, pH 7.3, 280 mOsmol. Excitability I/V curves (-20-180 pA, 10 pA steps, 500 ms) were measured in I-Clamp mode after gaining whole cell access. All data were analyzed offline using Clampfit (Molecular Devices). Excitability was measured as the total number of events during each current step.

*Behavioral health battery*

To screen for behavioral effects from systemic AAV.eB expression from experimental viruses the following battery of behavioral tests were performed in chronological order on the same mice:

*Open field testing:*

Mice were placed in a large circular arena (120 cm diameter) in dim light and activity was recorded for a period of 10 minutes. Data were analyzed using Ethovision software, where time in center, time on edge, and total distance were calculated.

*Elevated Plus maze (EPM):*

The EPM is plus-shaped arena in which two maze arms are sheltered by enclosed walls, and two arms are open platforms. Each arm length is 15 cm, with a center area of 49 cm^2^. Activity was recorded for a period of 10 min and analyzed using Ethovision software.

*Locomotor activity:*

Baseline locomotion was measured using locomotion chambers (Columbus instruments) that use infrared beam breaks to calculate ambulatory activity. Mice were singly housed in Allentown cages with reduced corncob bedding and provided with *ad libitum* access to food and water. Locomotion was monitored continuously over the course of 3 nights and 2 days.

**Statistics**

Raw data provided throughout the manuscript represent mean ± S.E.M. values. With the exception of voxel statistical tests (performed in ClearMAP software^13^), all other statistical analyses and data graphing was performed with Prism 10 software (GraphPad). Alpha levels for statistical significance were set at 0.05 for p and q-values. Probability levels of statistical significance from group mean comparisons are indicated in figures with an asterisk labeling convention of: *p<0.05; **p<0.01; **p<0.01; ****p<0.001. Two group comparisons were analyzed using two-tailed, unpaired Student’s t-test. One- or two-way ANOVA analysis of main effects were used to compare group means with one of two independent variables, respectively. Posthoc tests were chosen on an experiment-by-experiment basis and indicated in each figure legend. Region-by-region analysis of whole-brain datasets were performed using FDR-corrected multiple t-tests by the two-stage step-up method of Benjamini, Krieger, and Yekutieli^83^ on uncollapsed levels of all subregions from major brain ontologies.

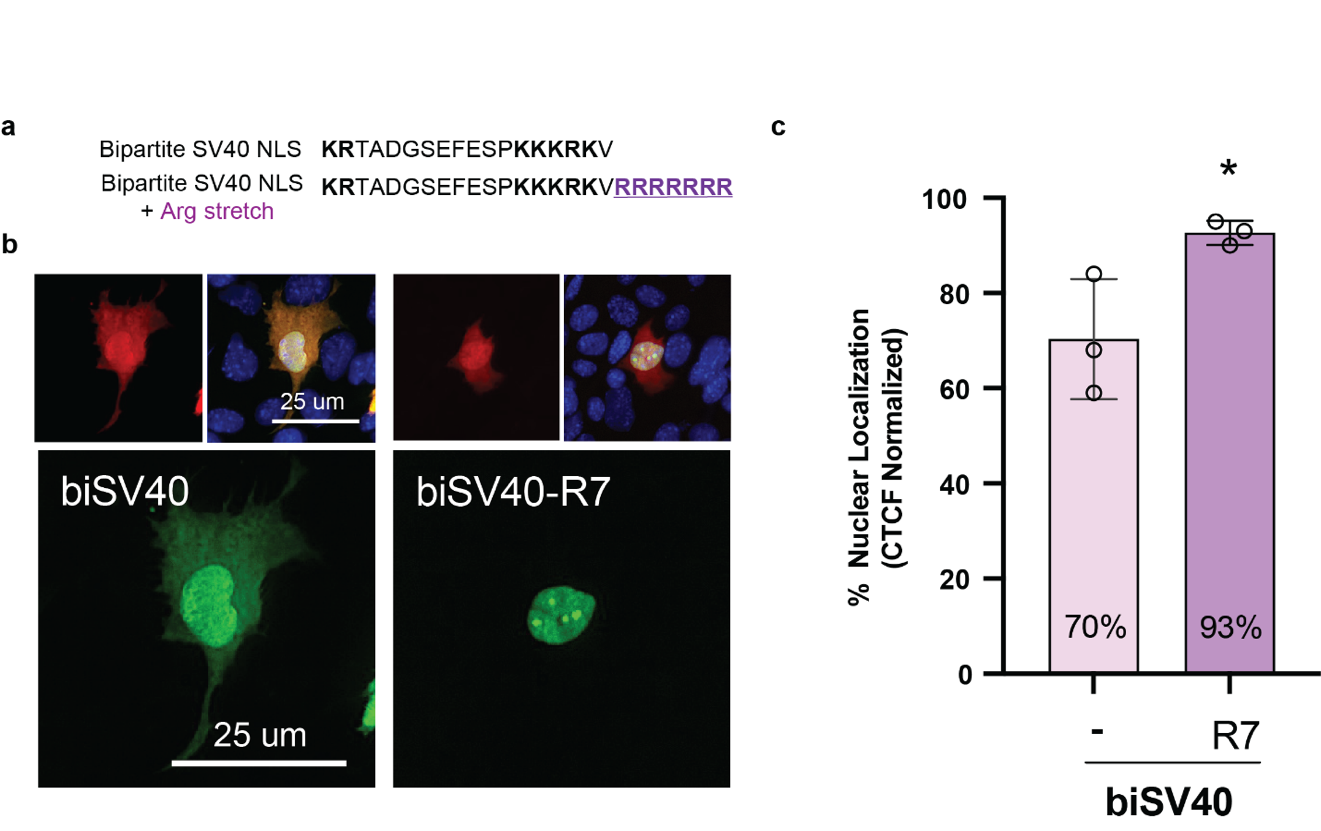

**Figure S1. Poly-R7 synergizes bipartite SV40 NLS activity *in vitro*. a,** Amino acid sequences of biSV40nls and biSV40nls-R7 tags. Basic amino acid residues are highlighted in bold **b,** Representative images of transient co-transfections with untagged mKate2 and biSV40- or biSV40-R7-EGFP-expressing constructs in an immortalized CLU198 embryonic hippocampal mouse cell line. **c,** Normalized corrected total cellular fluorescence (CTCF) of poly-R7 addition with biSV40nls (n=mean of 5 cells/3 cultures). Student’s t-test. **p<0.01.

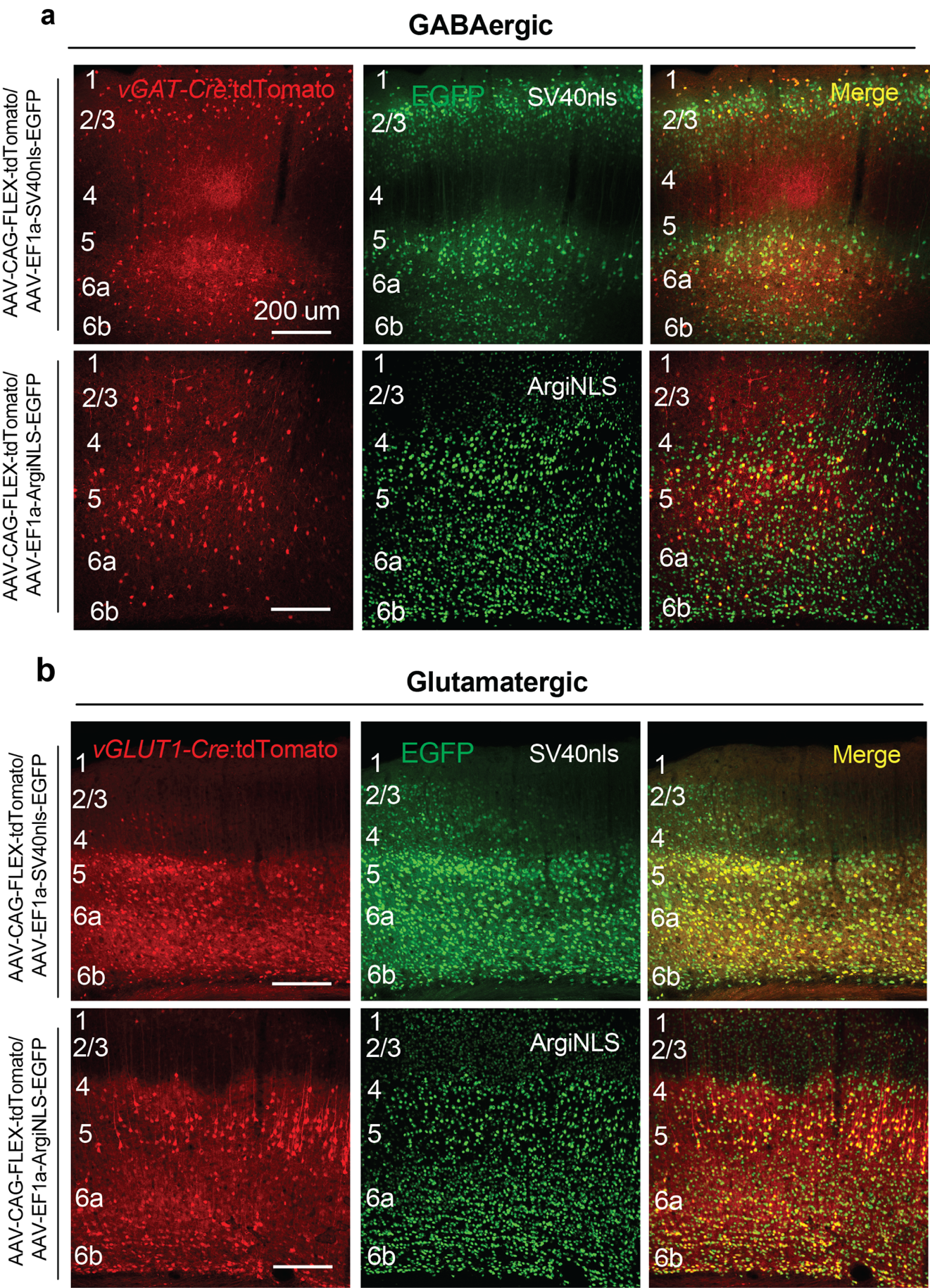

**Figure S2. Representative images of viral expression from local barrel cortex co-injections. a-b,** Example viral expression observed in a (**a**) *vGAT-Cre* or (**b**) *vGlut1-Cre* animal for Cre-dependent tdTomato and SV40nls- (top row) or ArgiNLS-EGFP (bottom row) viruses.

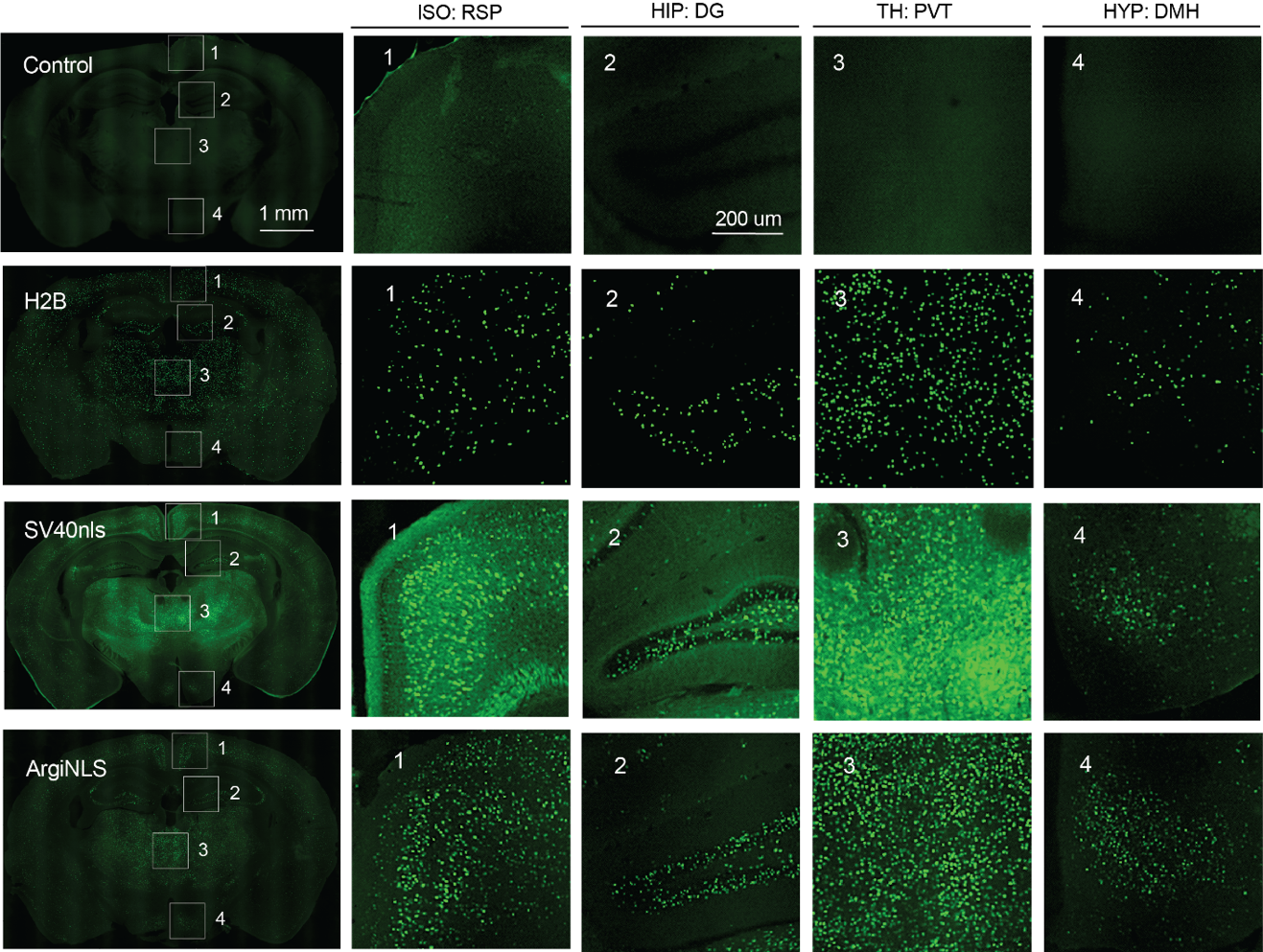

**Figure S3. Representative systemic PHP.eB viral EGFP expression from behavioral cohort.** Left: Example montage of coronal brain slices (~-1.94 Bregma) from representative samples from each group. Numbered inset FOVs are magnified to the right. Acronyms: HIP = hippocampus; DG = dentate gyrus; TH = thalamus; PVT = paraventricular thalamus; HYP = hypothalamus; DMH = dorsomedial hypothalamus; ISO = isocortex; RSP = retrosplenial cortex

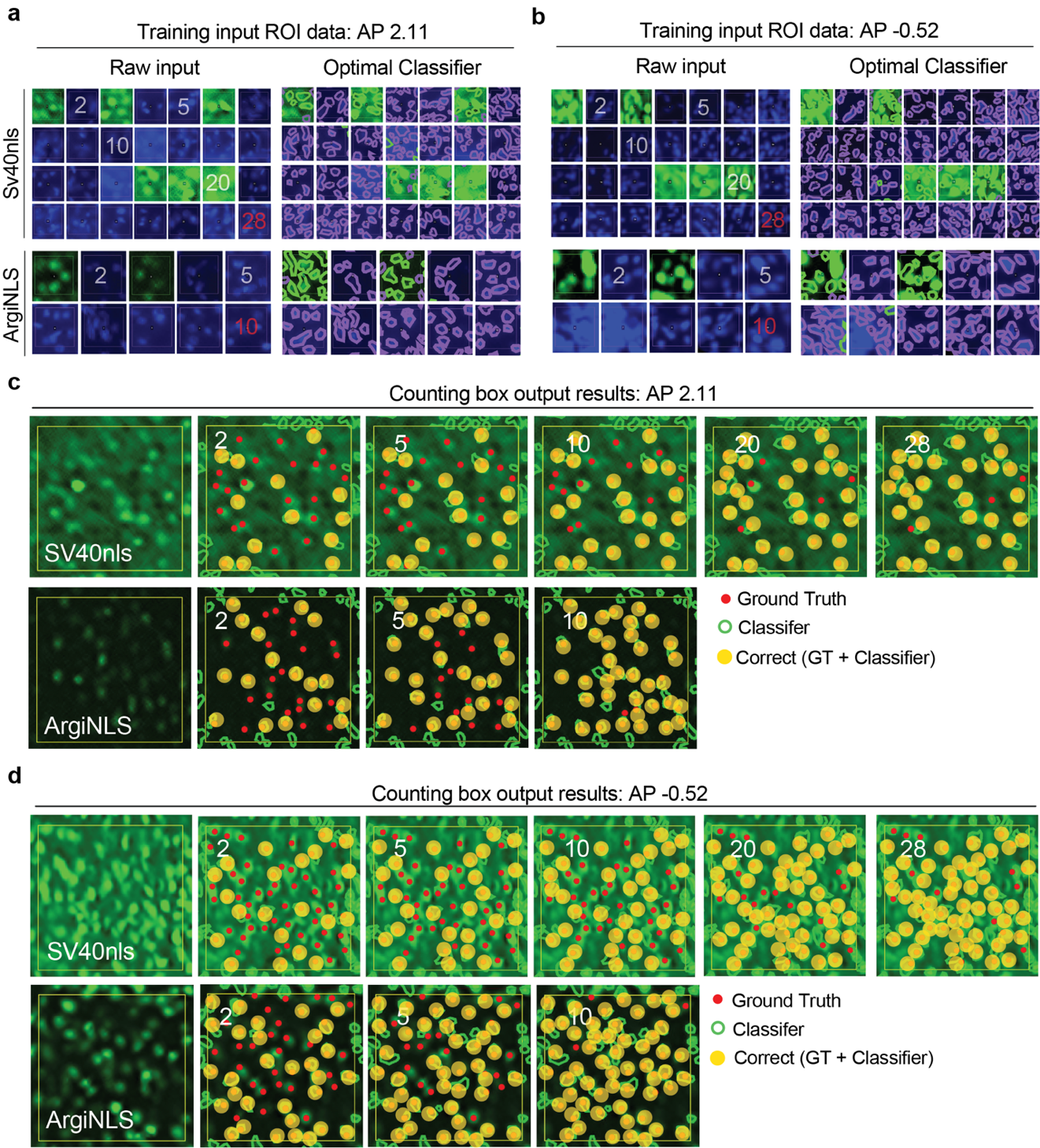

**Figure S4. Representative 2D raw images and performance of single-cell ML classification. a-b,** Classifier training input images for SV40nls (top) of ArgiNLS (bottom) from coronal brain section position corresponding to APs (**a**) 2.11 or (**b**) -0.52 from bregma. Raw input images are show to the left and the result shown to the right. The final training input image is numbered in red, white numbers indicate training input level. **c**-**d,** Raw images of counting box FOVs used for ground truth markup and comparisons with ML classifier at APs (**c**) 2.11 and (**d**) -0.52 for SV40nls (top) and ArgiNLS (bottom). White numbers indicate amount of input images (from **a-b**) used for classifier training.

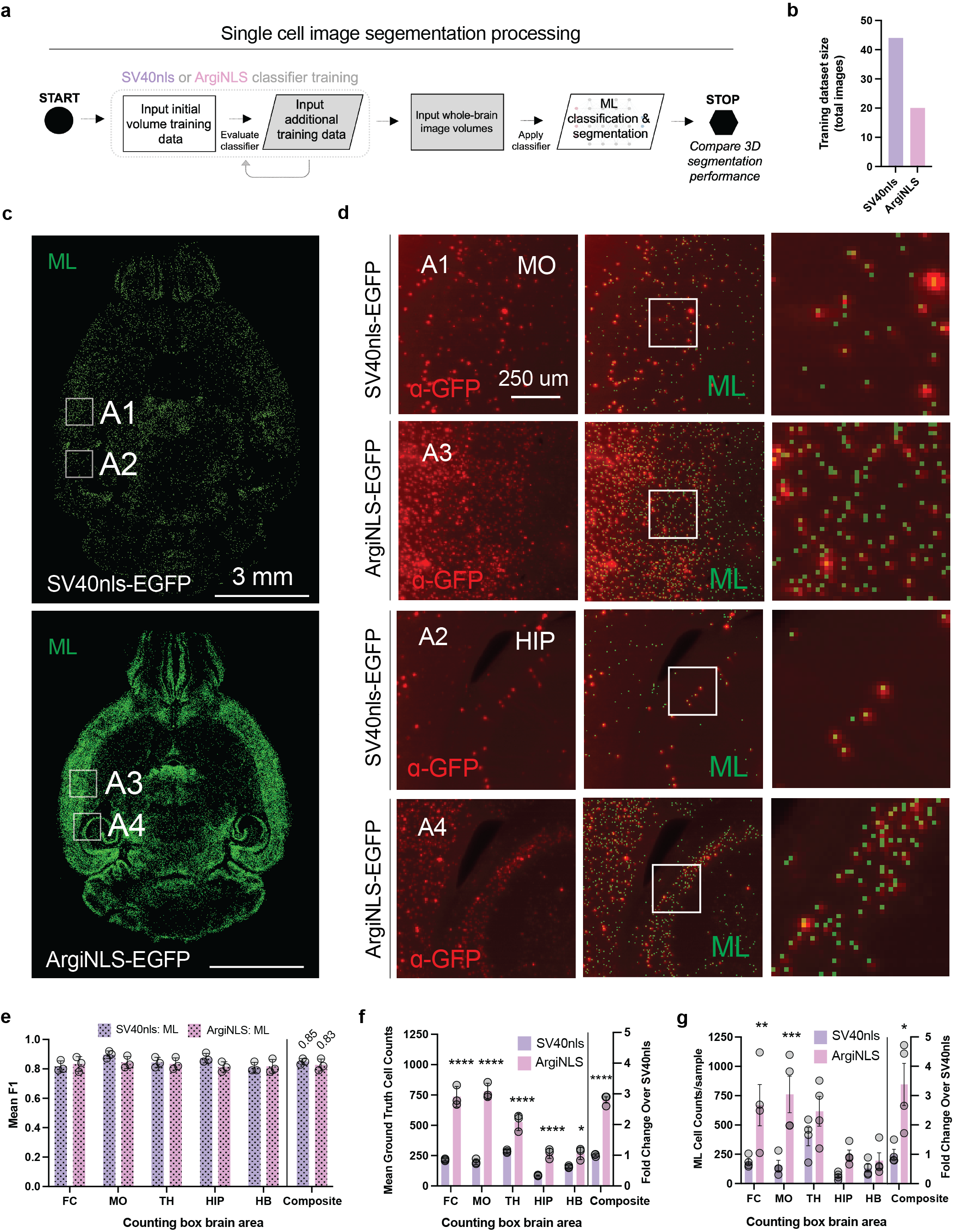

**Figure S5. Validation of ML-based brain-wide segmentation of single-cells.** **a,** Workflow of ML-based classifier training and application. **b**, Training dataset size of classifiers used in whole-brain classification. **c**, Representative images of ML-based centroids of segmentation from **Fig. 5b**, with additional insets of crop squares highlighting example areas (MO and HIP) used for classifier metric calculations. **d,** Magnified images of cropped insets from **c**, showing raw image (left; pseudo-colored in red), raw image with overlaid ML classification (middle), and zoomed inset of overlaid image (right). **e**, Mean F-score performance for each classifier across counting box brain areas. Composite F-score (see methods) is plotted and listed to the right. **f**, Comparison of mean (n = 3 samples) ground truth cell counts across counting box brain areas per expert. Composite fold change over SV40nls ground truth counts from all areas is plotted to the right. **g**, Comparison of ML counts across counting box brain areas for each sample imaged (n=4). Composite fold change over SV40nls ML cell counts from all areas is plotted to the right. The data in **e-f** were statistically analyzed using a 2-way ANOVA with Bonferroni-corrected posthoc test. Composite values were statistically analyzed with an unpaired t test. p<0.01; **p<0.05; **p<0.01; ****p<0.001. Acronyms: FC = frontal cortex; MO = primary motor cortex; TH = thalamus; HIP = hippocampus; HB = hindbrain

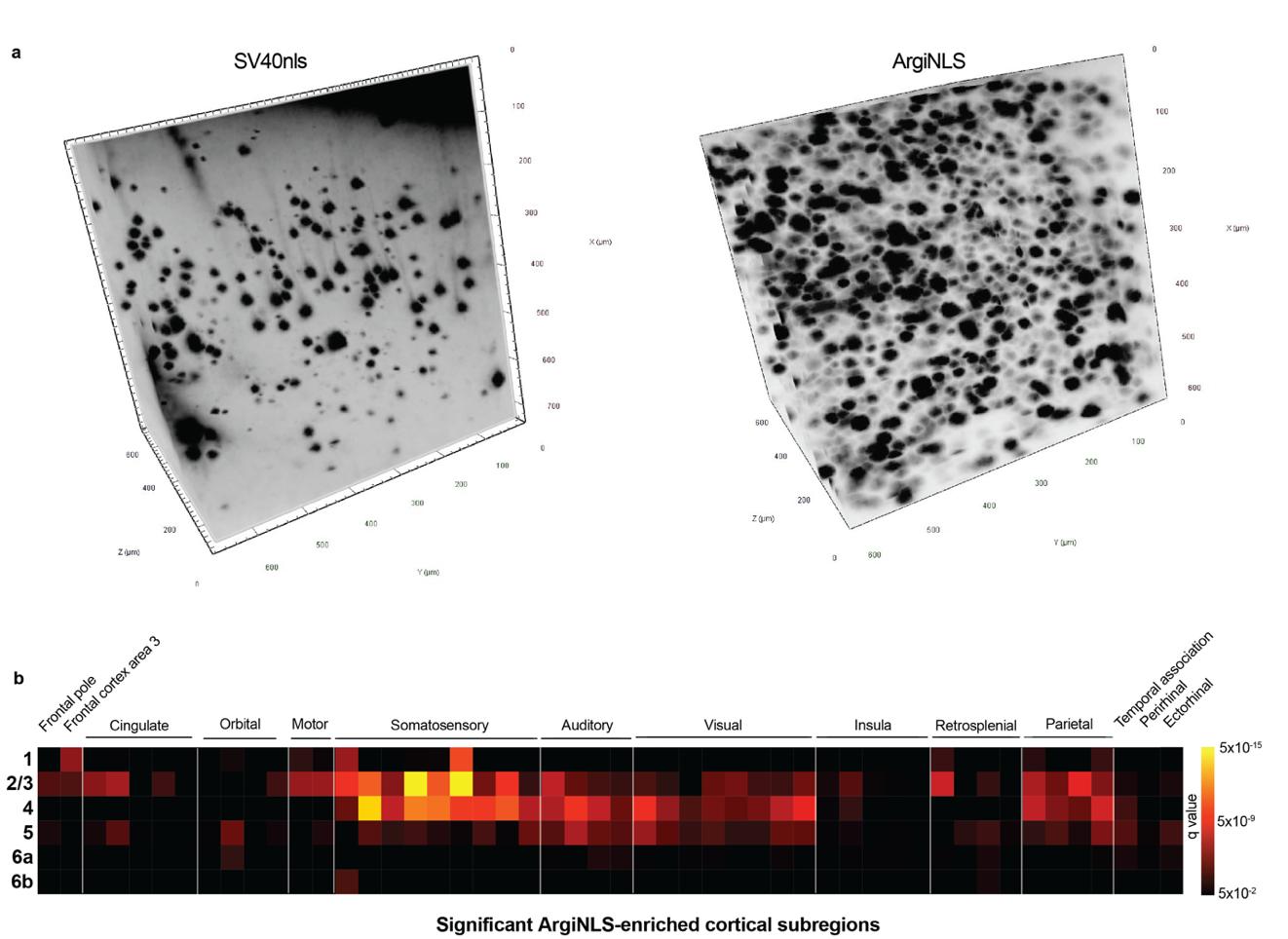

**Figure S6. Cortical differences in single-cell labeling and segmentation. a,** Representative raw volumetric 3D image crops displaying EGFP-stained single-cell labeling content in a 600 um^3^ cube located in the barrel cortex. **b,** Cortical heatmap of statistically significant differences from **Fig. 5f,** replotted across all cortical areas by layer.

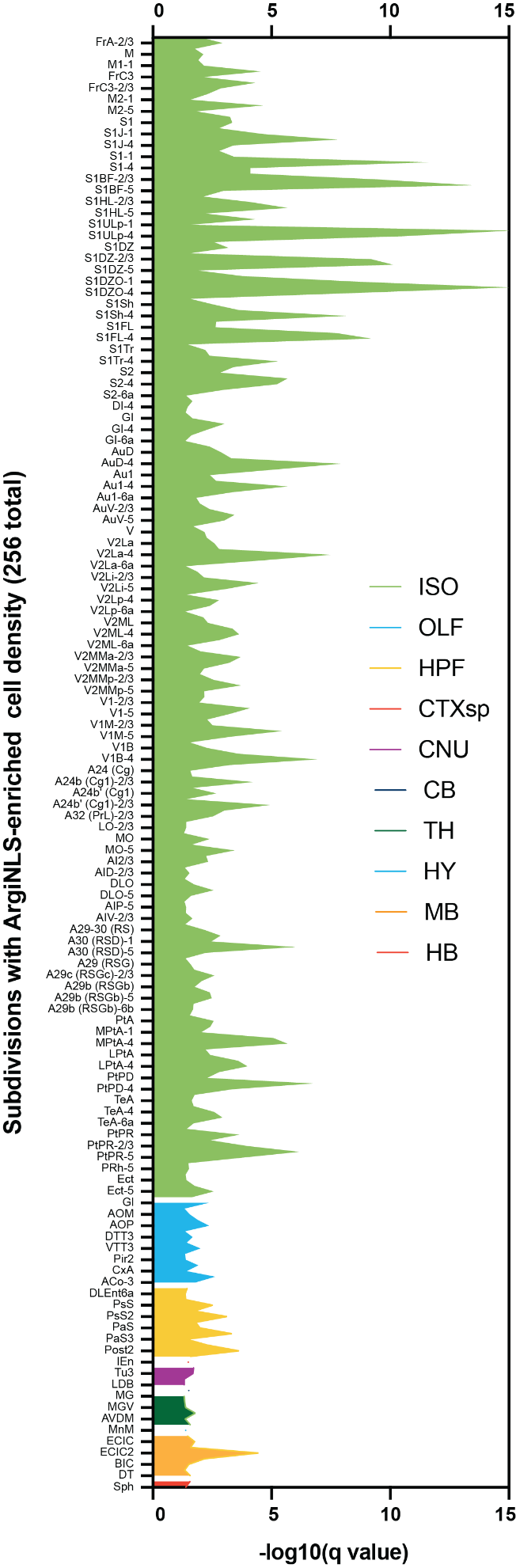

**Figure S7. ArgiNLS-enriched subregions. a,** Filled line graph display of -log10 q-values from all significant ArgiNLS-enriched subregions per major brain area. For legibility purposes, only every other subregion is listed. Full list and unabbreviated names are found in Supplementary Table 2.

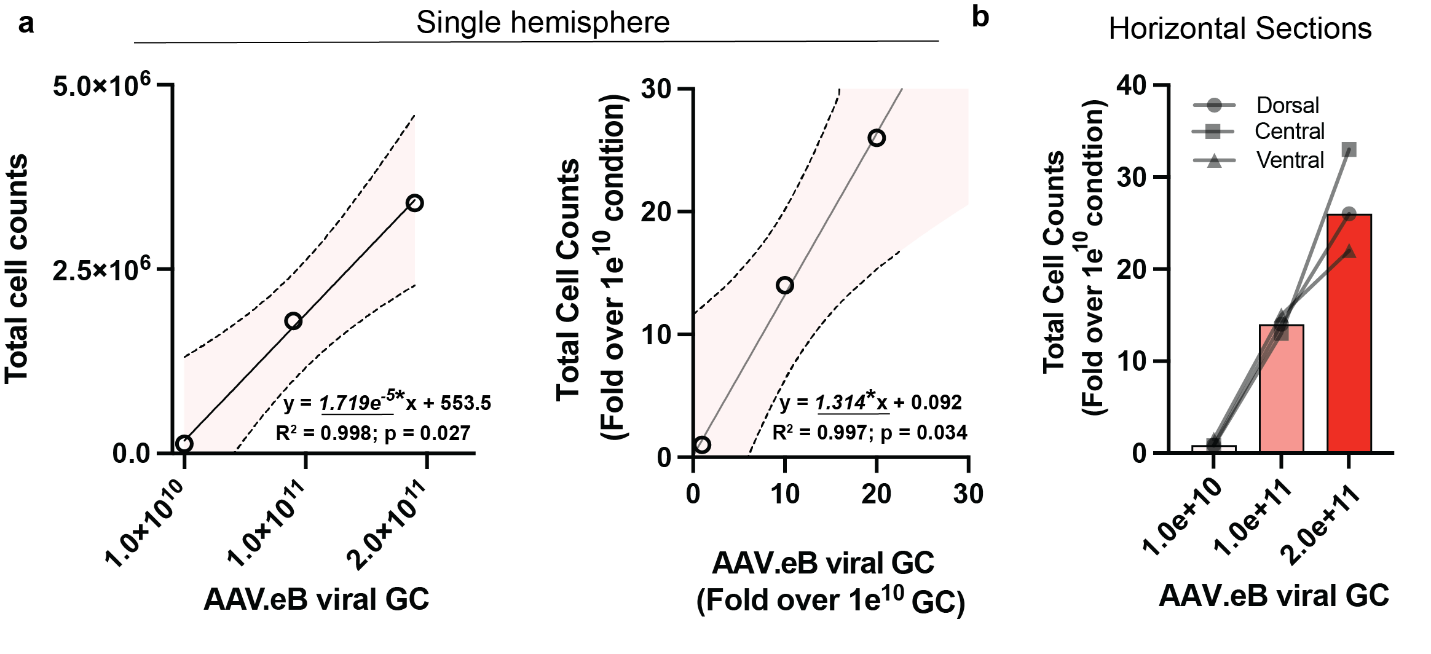

**Figure S8. Volumetric quantification of single-cells from systemic PHP.eB ArgiNLS-oScarlet labeling with 3 genetic payload amounts. a,** Left: Linear regression of total hemisphere cell counts versus AAV.eB viral GC payload amount. Right: Linear regression of fold-expressed cell counts versus fold-expressed GC payload. **b,** Bar graph of total cell counts from horizontal sections corresponding to 3 anatomical locations, taken from whole-brain samples.

| **NLS tag** | **Origin NCBI accession number** | **Sequence** | **Length** | ***pI*** | ***z*** | **CDS** |
| --- | --- | --- | --- | --- | --- | --- |
| SV40 | NP_0431271 | PKKKRKV | 7 | 11.5 | +4.9 | CCTAAGAAAAAGAGGAAGGTG |
| SV40-R3 | na | PKKKRKVRRR | 10 | 12.8 | +7.9 | CCTAAGAAAAAGCGCAAGGTGAGAAGGCGG |
| SV40-R5 | na | PKKKRKVRRRRR | 12 | 13.1 | +9.9 | CCTAAGAAAAAGCGCAAGGTGAGAAGGCGGCGGCGC |
| SV40-R7 | na | PKKKRKVRRRRRRR | 14 | 13.3 | +11.9 | CCTAAGAAAAAGCGCAAGGTGAGAAGGCGGCGGCGCCGTCGA |
| SV40-R9 | na | PKKKRKVRRRRRRRRR | 16 | 13.4 | +13.9 | CCTAAGAAAAAGCGCAAGGTGAGAAGGCGGCGGCGCCGTCGAAGGAGA |
| SV40-R11 | na | PKKKRKVRRRRRRRRRRR | 18 | 13.5 | +15.9 | CCTAAGAAAAAGCGCAAGGTGAGAAGGCGGCGGCGCCGTCGAAGGAGACGTCGC |
| biSV40 | NP_0431271 | KRTADGSEFESPKKRKV | 19 | 9.9 | +2.1 | AAAAGAACCGCCGACGGCTCCGAATTCGAGAGCCCTAAGAAAAAGCGCAAGGTG |
| biSV40-R7 | na | KRTADGSEFESPKKRKVRRRRRRR | 26 | 12.5 | +9.1 | AAACGCACCGCCGACGGCAGCGAGTTTGAGTCTCCCAAGAAAAAGCGCAAAGTAGAAAGGAGAAGAAGGAGAAGGAGGGGG |
| NPM2 | NP_001398932.1 | KRVAPQKQMSIAKKKKV | 17 | 11.6 | +6.1 | AAAAGGGTGGCTCCCCAGAAGCAGATGAGCATAGCAAAGAAAAAGAAGGTG |
| NPM2-R3 | na | KRVAPQKQMSIAKKKKVRRR | 20 | 12.8 | +9.1 | AAAAGGGTGGCTCCCCAGAAGCAGATGAGCATAGCAAAGAAAAAGAAGGTGAGAAGGCGG |
| NPM2-R5 | na | KRVAPQKQMSIAKKKKVRRRRR | 22 | 13.1 | +11.1 | AAAAGGGTGGCTCCCCAGAAGCAGATGAGCATAGCAAAGAAAAAGAAGGTGAGAAGGCGGCGGCG |
| NPM2-R7 | na | KRVAPQKQMSIAKKKKVRRRRRRR | 24 | 13.3 | +13.1 | AAAAGGGTGGCTCCCCAGAAGCAGATGAGCATAGCAAAGAAAAAGAAGGTGAGAAGGCGGCGGCGCCGTCGA |
| NPM2-R9 | na | KRVAPQKQMSIAKKKKVRRRRRRRRR | 26 | 13.4 | +15.1 | AAAAGGGTGGCTCCCCAGAAGCAGATGAGCATAGCAAAGAAAAAGAAGGTGAGAAGGCGGCGGCGCCGTCGAAGGAGA |
| NPM2-R11 | na | KRVAPQKQMSIAKKKKVRRRRRRRRRRR | 28 | 13.5 | +17.1 | AAAAGGGTGGCTCCCCAGAAGCAGATGAGCATAGCAAAGAAAAAGAAGGTGAGAAGGCGGCGGCGCCGTCGAAGGAGACGTCGC |

**Table S1. Sequence information of NLS tags tested.**

| **Construct** | **Promoter** | **FP Gene** | **Ex/Em** | **Condition** | **Addgene ID** |
| --- | --- | --- | --- | --- | --- |
| AAV-EF1ɑ-ArgiNLS-EGFP | EF1ɑ | EGFP | 488/507 | Constitutive | 211472 |
| AAV-EF1ɑ-ArgiNLS-AausFP1 | EF1ɑ | AausFP1 | 504/510 | Constitutive | 211473 |
| AAV-EF1ɑ-ArgiNLS-mGreenLantern | EF1ɑ | mGreenLantern | 503/514 | Constitutive | 211474 |
| AAV-EF1ɑ-ArgiNLS-mVenus-Q69M (ME) | EF1ɑ | mVenus ME | 515/528 | Constitutive | 211475 |
| AAV-EF1ɑ-ArgiNLS-oScarlet | EF1ɑ | oScarlet | 569/594 | Constitutive | 211476 |
| AAV-EF1ɑ-ArgiNLS-mKate2 | EF1ɑ | mKate2 | 588/633 | Constitutive | 211477 |
| AAV-EF1ɑ-ArgiNLS-miRFP670 | EF1ɑ | miRFP670 | 642/670 | Constitutive | 211478 |
| AAV-CAG-FLEX-ArgiNLS-EGFP | CAG | EGFP | 488/507 | Cre-dependent | 211479 |
| AAV-CAG-FLEX-ArgiNLS-AausFP1 | CAG | AausFP1 | 504/510 | Cre-dependent | 211480 |
| AAV-CAG-FLEX-ArgiNLS-oScarlet | CAG | oScarlet | 569/594 | Cre-dependent | 211481 |
| AAV-CAG-FLEX-ArgiNLS-mKate2 | CAG | mKate2 | 588/633 | Cre-dependent | 211482 |
| AAV-CAG-FLEX-ArgiNLS-miRFP670 | CAG | miRFP670 | 642/670 | Cre-dependent | 211483 |
| AAV-CAG-fDIO-ArgiNLS-AausFP1 | CAG | AausFP1 | 504/510 | Flp-dependent | 211484 |
| AAV-CAG-fDIO-ArgiNLS-oScarlet | CAG | oScarlet | 569/594 | Flp-dependent | 211485 |
| AAV-CAG-fDIO-ArgiNLS-mKate2 | CAG | mKate2 | 588/633 | Flp-dependent | 211486 |

**Table S2. ArgiNLS-tagged viral vector toolkit for optimized cell counting**

Movie S1 (separate file). Image stack fly throughs of coronally-sliced brain-wide voxelized cell densities for SV40nls-EGFP and ArgiNLS-EGFP infected group means (from Fig. 5) and significantly greater voxels for ArgiNLS (p<0.05).

Movie S2 (separate file). Volumetric renderings of systemic ArgiNLS-oScarlet expression at 3 AAV.eB GC payloads (from Fig. 6) amongst all axes and several zoom levels across the whole-brain.

Dataset S1 (separate file). Brain-wide cell count, density, and statistical test results related to Fig. 5 dataset. Provided as an excel file.
